# Multiscale Metabolic Covariance Networks Uncover Stage-Specific Biomarker Signatures Across the Alzheimer’s Disease Continuum

**DOI:** 10.1101/2025.07.10.663714

**Authors:** Juan Antonio K. Chong Chie, Scott A. Persohn, Ravi S. Pandey, Olivia R. Simcox, Paul Salama, Paul R. Territo, Alzheimer’s Disease Neuroimaging Initiative

**Affiliations:** Stark Neuroscience Research Institute, Indiana University School of Medicine; The Jackson Laboratory for Genomic Medicine; Wood College of Osteopathic Medicine, Marian University; School of Electrical and Computer Engineering, Purdue University; Department of Medicine, Indiana University School of Medicine

**Author notes:** Data used in preparation of this article were obtained from the Alzheimer’s Disease Neuroimaging Initiative (ADNI) database (adni.loni.usc.edu). As such, the investigators within the ADNI contributed to the design and implementation of ADNI and/or provided data but did not participate in analysis or writing of this report. A complete listing of ADNI investigators can be found at: http://adni.loni.usc.edu/wp-content/uploads/how_to_apply/ADNI_Acknowledgement_List.pdf.

## Abstract

**Background:** Connectomics studies analyze neural connections and their roles in cognition and disease. Beyond regional comparisons, recent research has revealed inter-regional brain relationships via graph theory of brain network connectivity. Within these networks, path length measures a network’s efficiency in communication. These connections can be quantified as inter-subject covariance networks related to functional connectivity, with alterations reported in neurodegenerative diseases.

**Methods:** Retrospective analysis of ADNI ^18^F-FDG PET images using metabolic covariance analysis and hierarchical clustering was used to assess regional brain networks in subjects from cognitively normal (CN) to AD. We evaluated AD stage changes by calculating whole brain entropy, connection strength, and clustering coefficients. Additionally, estimates of shortest path for positive and negative correlations as a measure of network efficiency. We also developed a novel region set enrichment analysis (RSEA) to detect brain functional changes based on metabolic variations. Results were aligned with transcriptomic signatures and clinical cognitive assessments.

**Findings:** In AD subjects, whole brain metabolic connectivity revealed an increase in entropy, connection strength, and clustering coefficients, which indicates brain network reorganization as compensatory mechanisms of pathological disruption. As AD advances, path lengths between brain regions decrease from CN to MCI; however, path lengths significantly increased in AD. RSEA indicated functional changes in motor, memory, language, and cognition functions related to disease progression.

**Interpretation:** Metabolic covariance analysis of whole brain, and regional connectomics, track with AD progression. Moreover, path lengths permitted AD stages determination via alterations in brain connectivity. Furthermore, RSEA facilitated the identification of functional changes based on metabolic readouts.

**Funding:** NIH grant T32AG071444

## 1. BACKGROUND

Alzheimer’s disease (AD) and related dementia (RD) are a suite neurodegenerative disorder characterize by progressive cognitive decline, which currently affects 6·9 million people in the US and projected to grow to 13·8 million by 2060.^1,2^ Despite extensive research, the causes of ADRD remain unclear, emphasizing the need for continued investigation.^1^ Recent studies suggest that accumulation of amyloid-β plaque and hyperphosphorylated tau tangles leads to neuroinflammation, oxidative stress, metabolic dysregulation, synaptic disruption, and neuronal dysfunction, thus leading to cognitive decline, brain networks disruption, and neurodegeneration.^3^

Metabolic reprogramming theory^4,5^ suggests that energy defects during aging cause metabolic dysregulation in neurons and astrocytes, highlighting the interaction between oxidative phosphorylation and glycolysis.^4,5^ Importantly, this bioenergetic support dictates neuronal viability, while aging process determines the long-term efficiency of energy production.^4^ Under these conditions, the decline in enzyme function involved in metabolic regulation during aging leads to dysregulation, and ultimately neuronal degeneration.^4^ Moreover, the theory proposes that the associated increase in energy demand is compensated by the inverse Warburg effect (upregulation of oxidative phosphorylation activity in neurons) and the Warburg effect (upregulation of glycolysis in astrocytes).^4^ These two processes increase lactate production as a compensatory mechanism to provide an adequate energy source for neurons.^4^ Support for this premise comes from work in our lab, which has shown in mouse models containing human *APOE*^*e4*^ genes, that depending on genetics, age and sex, metabolism is dysregulated with respect to blood flow.^6^ Similarly, recent work from our lab in humans across a wide array of genetics, ages, sexes, and disease states show a striking similarity with AD mouse biology, thus helping to support this premise.^7^ The use of positron emission tomography (PET) of glycolytic metabolism has proven useful in predicting and diagnosing ADRD.^8^ Studies^9–11^ have shown that early prodromal stage of the disease involves hypermetabolism, while regional hypometabolism is observed in key brain areas at later stages of the disease.^12,13^ Additionally, recent studies using ^18^F-2-fluorodeoxyglucose (^18^F-FDG) as a metabolic imaging biomarker,^14^ showed that disease progression impacts brain connectivity networks,^15^ resulting in default mode network variations and AD-related patterns.^14^

Recent connectomics modeling studies^9,15^ have shown that brain network organization, structure, and function at various structural scales can identify alterations associated with neurological and psychiatric disorders^16^, and establish how neural connections contribute to cognitive functions and disease progression.^16^ Consequently, connectomics modeling has enhanced our understanding of brain networks by enabling the analysis of inter-regional relationships through graph theory and brain network connectivity beyond simple regional comparisons.^17^ Moreover, these relationships derived through connectomics modeling of surrogate metabolic imaging biomarkers are frequently referred to as metabolic connectivity networks^18^, and provide detailed insights into how brain regions coordinate activity to support complex behaviors. Preclinical and clinical studies^9,15,19,20^ on metabolic connectivity in neurodegenerative diseases, such as AD, have reported alterations in metabolic connectivity networks (e.g., whole brain entropy, connection strength, clustering coefficients) associated with functional connectivity changes, thereby supporting the brain reprogramming^4^ and energy failure^21^ theories.

Although brain connectomics has become a major research field in neuroscience, molecular imaging is not widely used due to the dominance of magnetic resonance imaging (MRI) in neuroimaging.^18^ Hence, current brain connectivity networks are mainly based on MRI modalities (i.e., BOLD, diffusion MRI, fMRI), which limits the understanding of inter-regional brain connectivity, since MRI is based on T2/T2* changes with alterations in blood volume as a surrogate for brain metabolism,^22^ and thus overlooks the complex biochemical processes and electrical signaling related to brain activity.^18^ Therefore, in this study, we explored metabolic connectomics changes across the AD spectrum, ranging from cognitively normal (CN) to AD, using ^18^F-FDG as a metabolic surrogate^14^ and network covariance analysis. To explore and quantify brain functional changes in these networks, we developed a new method known as region set enrichment analysis (RSEA) to examine these measures. Moreover, we extended existing network metrics, by computing functional path lengths, permitting detection of network remodeling and change in network efficiency over the disease spectrum. Based on this, we hypothesized that the pattern of the alterations in the metabolic connectivity networks throughout the disease spectrum can be characterized to determine inter-regional changes and their contribution to cognitive functional decline and disease staging, thus allowing sensitive and specific means to assess disease progression, patient stratification, and therapeutic response monitoring.

## 2. MATERIALS AND METHODS

### 2.1 Dataset

Data used in the preparation of this article were obtained from the Alzheimer’s Disease Neuroimaging Initiative (ADNI) database (adni.loni.usc.edu). The ADNI was launched in 2003 as a public-private partnership, led by Principal Investigator Michael W. Weiner, MD. The primary goal of ADNI has been to test whether serial MRI, PET, other biological markers, and clinical and neuropsychological assessment can be combined to measure the progression of mild cognitive impairment (MCI) and early Alzheimer’s disease (AD).^23^

Specifically, we mined data from ADNI phases 2 and 3 for subjects scanned with ^18^F-FDG PET and clinical cognitive assessment (CCA) results for Alzheimer’s Disease Assessment Scale-Cognitive Subscale (ADAS-Cog), Clinical Dementia Rating (CDR), and Montreal Cognitive Assessment (MoCA) within 180 days. A total of 431 cases were obtained, which retained the following split: 98 CN, 76 early MCI (EMCI), 129 MCI, 42 late MCI (LMCI), and 86 AD. The age range of the subjects was from 55 years to 95 years with an average age of 73·60±7·44 years, and the sex split between the cases were 55% for males and 45% for females. In addition, vitals (age, height, weight) and APOE genotypes for these subjects were obtained. The APOE distribution was 6·73% for APOE2 (ε2/ε2 and ε2/ε3), 61·48% for APOE3 homozygotes (ε3/ε3), and 31·79% for APOE4 (ε2/ε4, ε3/ε4 and ε4/ε4). For complete demographics, see Table S1.

### 2.2 Image preprocessing

To evaluate regional cerebral glycolysis, glucose uptake was measured via ^18^F-FDG PET as surrogate biomarker readout for glycolytic metabolism.^14^ PET images were acquired and reconstructed using standard parameters (see Table S2) developed by ADNI^23^. To permit direct comparison in a common reference space (i.e. Montreal Neurological Institute + Harvard-Oxford Cortical + subcortical(RRID: SCR_001476) + FSL Probabilistic Cerebellar Atlases; MNI152+), all images were co-registered using a 3D deformable registration based on total variation developed by our lab.^24^ For a list of brain regions and indices, see Table S3.

Post-registration, mean intensities for 59 unilateral brain regions were extracted via the MNI152+ atlas for the ^18^F-FDG images. Typically, regional analysis utilizes standardized uptake values (SUV), assuming body mass may serve as a reasonable proxy for the tracers volume distribution; however, this assumption cannot be met for all subjects in an aging study whose body mass and composition are non-uniform.^25,26^ To overcome these limitations, SUV ratios (SUVR) have been used, which represent tracer uptake as a ratio of the region/volume of interest to a reference structure (e.g., cerebellum, whole brain).^27^ In the case of ADRD studies, where disease progression is non-uniform throughout the brain, the selection of a reference region may bias the interpretations;^28^ therefore, to eliminate this bias, SUVRs have been computed relative to whole brain^27^. In the current study, SUVRs were computed as follows:

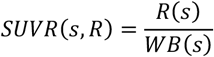

where, *s, R*, and *WB* denote the subject, mean value of the region/volume of interest for subject *s*, and whole brain mean value for subject *s*, respectively. For each regional SUVR, values were converted to z-scores relative to the CN population as follows:

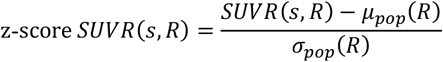

where, *s, R, μ*_*pop*_, and *σ*_*pop*_ are the subject, region/volume of interest, mean SUVR of the reference population, and standard deviation of the reference population. The z-score SUVR values are used to calculate and determine the significance of each region across the disease stages using Student’s t-test with p<0·05.

### 2.3 Metabolic Connectomics Analysis

Metabolic covariance analysis was conducted in MATLAB 2024b to evaluate whole brain and sub-network changes in the metabolic covariance networks. The following open-source packages were used: Brain Connectivity Toolbox (BCT),^29^ generalized Louvain modularity,^30^ hierarchical consensus clustering,^31^ and CovNet developed by our lab.^9^

Following our previous preclinical work, metabolic covariance networks were computed for 5 disease stage groups (CN, EMCI, MCI, LMCI, AD) across subjects within the group. The resulting covariance network matrix for each group resulted in a region-by-region (59×59 region) adjacency matrix that indicates the degree of similarity of regional ^18^F-FDG readout among all region pairs. Two covariance network variants were generated: weighted unthresholded networks (WUN) and weighted thresholded networks (WTN) at p<0·05 correlation significance using CovNet.^9^

Using the Brain Connectivity Toolbox,^29^ the following network measures were computed on WTN for each disease stage: network density (number of connections), node degree (number of connections for each region), positive and negative nodal strengths (sum of the positive/negative covariance values for each region), and clustering coefficient (number of fully connected triangles around a node based on the weight and sign of the connections).^32^

Community detection, or modularity maximization, is the process of identifying groups of nodes in a network with greater intra-group connectivity than inter-group connectivity. In this study, this process is performed using multiresolution consensus clustering (MRCC),^31^ which identifies communities across spatial scales and quantitatively determines a consensus partition from the set. MRCC was applied to the WUN as they contain the most information. These derived consensus partitions were used to investigate differences between the modules obtained for each group, based on their metabolic SUVR values, modules obtained from the CN group for males and females, and were used to analyze disease progression.

### 2.4 Metabolic Shortest Path Analysis

WTN were separated into two adjacency matrices, one for each: positive and negative correlations. Under the assumption that these matrices can be viewed as undirected graphs, using the absolute values of the entries of these matrices, we obtained the positive and negative shortest path trees between all pair-wise combinations of regions using Dijkstra’s algorithm. Although shortest path analysis has been reserved for structural connectomics, which is analogous to measuring the shortest distance between nodal pairs,^16^ this measure is thought to represent most efficient routes of information exchange between brain regions,^33,34^ and is analogous to the length of the wires connecting nodes in a network. However, in the context of metabolic networks, this measure represents metabolic interactions that minimizes the number of functional steps to link nodal pairs, and elucidates how metabolic processes disturbances propagate through the network,^35^ and therefore more analogous to the voltage across a given wire. As such, the shortest paths between regions were ordered into a 59×59 metabolic distance matrix (MDM), where the upper triangular part (UMDM) represents the positive path lengths between regions and the lower triangular part (LMDM) represents the negative path lengths between regions. This can be described by the following:

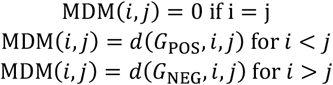

where *i* indexes *i*^th^ the brain region, *j* the *j*^th^ brain region, *MDM* the entry (*i, j*) of the MDM, *G*_POS_ the undirected graph using the positive values of the WTN, *G*_NEG_ the undirected graph using the absolute value of the negative values of the WTN, and *d* is the shortest distance between region *i* and region *j* for graph *G* using Dijkstra’s algorithm. In addition, to understand the network balance between positive and negative pathways, differences between positive and negative shortest-path trees were computed. An inter-regional path length comparison was performed to determine the difference between disease stage groups to evaluate the effect of disease progression in path lengths.

### 2.5 Region Set Enrichment Analysis

Region set enrichment analysis (RSEA) was developed to detect and quantify brain function changes based on metabolic variations. It is worth mentioning that although we developed this method in the context of metabolic alterations, it can be translated to other readouts. This method weighs changes in each region and relates them to functional changes, that are split into 10 functional-set-enrichment categories (FSEC): auditory, autonomics, cognition, emotions, language, memory, motor, sensorial, speech, and visual. A 59×10 matrix is generated, whose the columns represent the FSEC, and rows the brain regions. For each brain region, a column is assigned the value of 1 if the brain region is related to the relevant FSEC; otherwise, it is assigned a value of 0. (The list of functions by region plus literature references used to generate this matrix is found in Table S4.) Then, each value is normalized using the following:

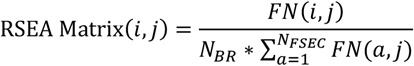

where, *i* indexes the brain region, *j* the FSEC, *N*_*BR*_ the number of brain regions, *N*_*FSEC*_ the number of FSEC, and *FN*(*i, j*) the (*i, j*) entry of the RSEA matrix. To perform RSEA analysis, differences between SURV values of each brain regions relative to a reference group are multiplied elementwise across its corresponding RSEA matrix row. RSEA can be performed across multiple modalities: whole brain, region clustered by significance, and regions clustered by connectomics modules.

For the whole brain, a single score value for each FSEC was obtained by summing each column of the RSEA matrix. For regions clustered by significance, p-values using Student t-tests for each brain regions relative to CN was calculated for each disease stage. Regions were then divided into 4 quartiles. The smallest p-value of each brain region for all stages was chosen, and regions were sorted from smallest to largest p-value. Subsequently, the 59 brain regions were divided into quartiles: the first quartile included regions 1 through 15, the second quartile included regions 16 to 30, the third quartile included regions 31 to 45, and the fourth quartile included regions 46 to 59. For each quartile, a single score value for each FSEC was computed by summing the columns of each region. Similarly, for regions clustered by connectomics modules, the MRCC modules computed for the CN group were used to cluster the regions. The FSEC score was generated by summing the columns of the regions within the corresponding module.

### 2.6 Transcriptomics Analysis

The WGCNA algorithm^36^ was employed to identify gene co-expression network, across the blood samples, utilizing the following parameters: soft threshold power = 6, minimum module size = 30, merge cut height = 0·25. Key genes (hub genes) were identified from each module using module eigengene (ME). In addition, Pearson correlations between module eigengene and participant characteristics, including sex, ^18^F-FDG, and disease stage were calculated. These module-trait associations are visualized via the ComplexHeatmap package in R. For detailed WGCNA procedures, refer to the Supplemental Methods.

Functional annotations and enrichment analysis were performed using the Bioconductor package clusterProfiler^37^ in R. Gene ontology (GO) terms were enriched using enrichGO, while KEGG pathway enrichment analysis was performed using enrichKEGG. Enriched functional categories within gene modules were compared using compareCluster. A significance threshold of 0·05 was set for all enrichment analyses, utilizing Benjamini-Hochberg adjusted p-values.

The biological domains by Cary et al.^38^ were used to perform functional enrichment analysis relative to AD to obtain AD-associated endophenotypes. Subsequently, we conducted Gene Set Enrichment Analysis (GSEA)^39^ and categorized the results into biological domains and sub-domains based on the GO ID of enriched terms via the gseGO function from the clusterProfiler R package.^37^ For further details, refer to the Supplemental Methods.

### 2.7 Statistical Significance

Statistical analysis of significant brain regions was evaluated using Student’s t-test (p<0·05) on the z-scored SUVR values relative to CN. The correlation between nodal measures, path metrics, and CCA were aligned to evaluate their relationship across the disease spectrum, using Pearson’s correlation and Student’s t-test (p<0·05). Nodal measures between groups were compared via 2-sample Kolmogorov-Smirnov (KS) tests and RSEA modules significance was determined via Student’s t-test.

## 3. RESULTS

3D image registration for all ^18^F-FDG images was the evaluated using structural similarity index measure (SSIM), to assess similarity in structural features between images, and their registered version. Quality assurance was performed for all cases by a team of experts to ensure proper alignment and scan coverage. The average SSIM was 0·91±0·04 for (N=431), indicating a high degree of alignment quality.

### 3.1 Metabolic Covariance Network Analysis

Covariance and connectomics analysis were used to assess network-level metabolic changes. Figure S1 shows the unthresholded metabolic covariance matrices across the disease spectrum, with brain region names in Table S3. As the disease progresses, the WTN become sparse and fragmented, especially in females as shown in Figure S2.

Network densities (Figure 1·A) measure the number of connections, with a density of 0 indicating no connections between nodes, and 1 indicating a fully connected network (every node is connected to every other node in the network). At p<0·05, male node density oscillates between 0·3068 and 0·4664, while female node density ranges from 0·3010 to 0·3904. Both sexes show a ‘W’ pattern where the drop in node density occurs during transition stages (EMCI/LMCI). Comparing between CN, MCI, and AD, male node density increases as disease progresses, whereas female node density remains relatively constant.

**Figure 1.**
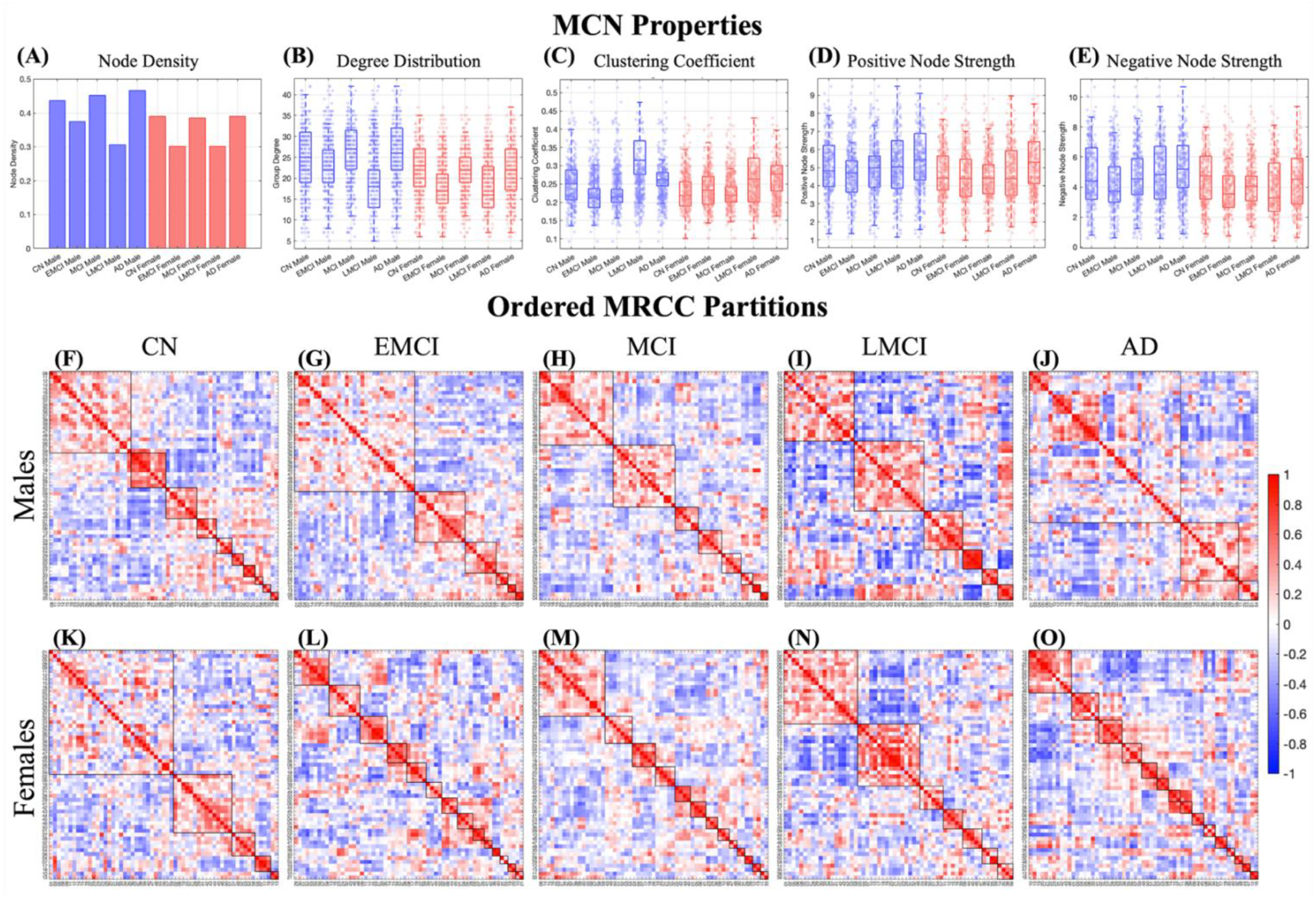
Properties of Thresholded Metabolic Covariance Networks and Ordered MRCC Partitions Across the Disease Spectrum. The following metrics were computed using thresholded metabolic covariance networks (MCN) with p<0·05: (**A**) Network densities, (**B**) Degree distribution, (**C**) Clustering Coefficient, (**D**) Positive Nodal Strength, and (**E**) Negative Nodal Strength. These metrics show variations across the disease spectrum. Notably, while both sexes exhibit similar changes, when compared to the same disease stage, each sex displays alterations to a different extent. This suggests that the brain networks restructure at different rates, thus exhibiting dimorphism. (**F-J**) Ordered MRCC partitions for males. The number of modules varies across different disease stages: 10 for CN, 6 for EMCI, 8 for MCI, 6 for LMCI, and 3 for AD. This suggests that brain regions reorganize as AD progresses. In male cases, the network begins consolidating which may be due to reduced functionality or a shift in metabolic demand priorities as an attempt to preserve balance in the brain. (**K-O**) Ordered MRCC partitions for females. The number of modules varies across disease stages: 5 for CN, 14 for EMCI, 13 for MCI, 7 for LMCI, and 15 for AD. Females exhibit an increase in the number of modules, indicating a sex dimorphism in network clustering compared to males, who show a decrease in modules. However, this suggests that the brain reorganization places more stress on female networks, leading to fragmentation. Consequently, the restructuring results in more specialized modules.

Degree distributions (Figure 1·B) denote the frequency of connections for a node, and is used to detect network hubs. Greater number of highly connected nodes make the network more heterogeneous and centralized. At p<0·05, the degree distribution for both sexes also shows a ‘W’ pattern similar to node density. Both sexes experience a drop in the LMCI phase, suggesting reduced information exchange efficiency between brain regions.

The clustering coefficient (Figure 1·C) measures how closely nodes cluster together, with 0 indicating no clustering and 1 indicating full connectivity. Thus, this measure represents the probability that two nodes are linked to a third node, where a high clustering coefficient implies fewer long-range connections. At p<0·05, variations in clustering coefficients are observed across different diseases for both sexes, with higher clustering coefficients occurring at the late stages.

Positive and negative nodal strength (Figures 1·D-E) represent the sum of positive or negative nodal weights. These metrics are utilized to assess the connections to other nodes, where high strengths indicate more central and influential impact on nodes within the network. At p<0·05, changes in nodal strength are observed in both sexes across the disease spectrum, with both positive and negative strengths following a similar pattern. Table S5 shows the KS tests comparison of the network analysis measurements for all pairwise groups combinations.

### 3.2 Modularity Structure of Metabolic Covariance Networks

By employing a data-driven approach with MRCC, using 10,000 permutations, consensus communities (modules) were identified for each unthresholded metabolic covariance network. The number of modules for males/females (Figure 1·F-J/1·K-O) was as follows: 10/5 for CN, 6/14 for EMCI, 8/13 for MCI, 6/7 for LMCI, and 3/15 for AD. This indicates the presence of a sexual dimorphism in the clustering of the networks, as the number of modules decreases for males while they increase for females. The number of modules can be used to assess the coherence among regions within the network, with a higher number of modules indicating a more disordered or fragmented network. This result suggests that the brain undergoes restructuring of the modules in relation to disease progression to keep the balance in the brain.

### 3.3 Shortest Path Distances

Each entry (*i, j*) of the distance matrix (Figure S3) represents the shortest path length between regions for positive connections in the UMDM and negative connections in the LMDM. Values on the diagonal are set to 0 as they represent the path length of a region with itself. Qualitative analysis of these matrices across the disease spectrum indicates that the largest increase in distance occurs during the LMCI phase for both sexes.

Quantitative analysis of distance matrices was performed using CN (Figure 2) as a common reference across all stages, and the previous stage (Figure S4) to observe changes over disease progression. Both sexes showed similar patterns for the two references. Relative to CN: EMCI, LMCI, and AD had increased path lengths, while MCI had decreased path lengths. Relative to the previous stage: EMCI and LMCI had increased path lengths compared to their prior stages (CN and MCI), whereas MCI and AD had decreased path lengths compared to their prior stages (EMCI and LMCI). The varying path lengths across the disease spectrum suggest changes in energy expenditure and lowered efficiency.

**Figure 2.**
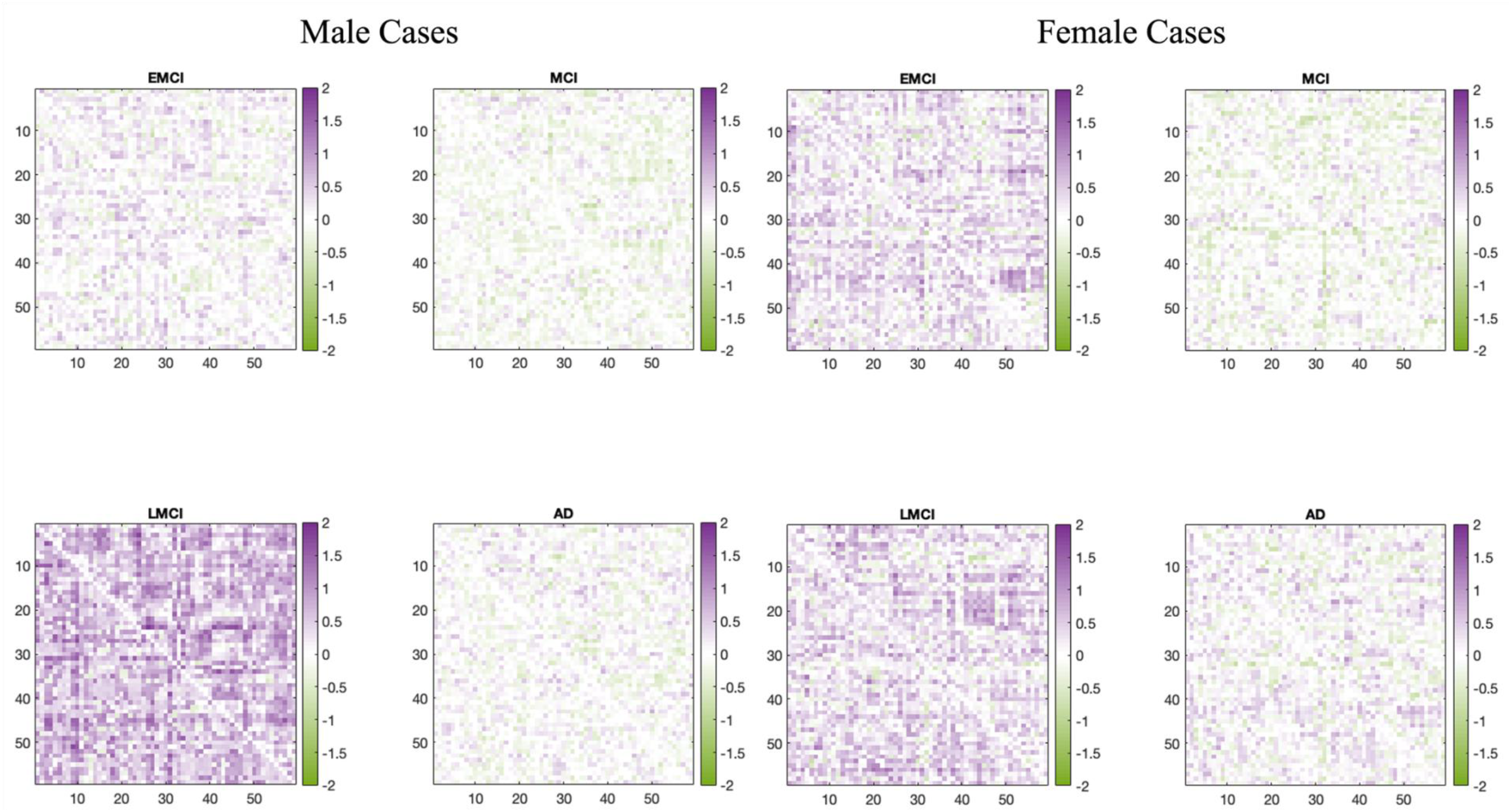
Comparisons of the Pathways Relative to CN. Quantitative analysis comparing CN to each other disease stage reveals that EMCI, LMCI, and AD have increased path lengths, while MCI has shown a decrease. Typically, an increase in path length indicates less efficient networks, while a decrease suggests more efficient networks. However, in this case, the MCI increase is attributed to the larger metabolic demand, which makes the network appear more efficient despite consuming more energy. Consequently, these changes in path lengths across the disease spectrum suggest alterations in energy expenditure and efficiency.

Table S6 shows the total path lengths, number of edges, and number of path segments for both positive and negative paths. These changes create an ‘M’ pattern, with the peak at LMCI aligning with connectomics analysis results, indicating that the largest change occurs at this stage. After normalizing path lengths by the number of segments, the difference falls within the interval [-0·0039, 0·0047]. This indicates that the magnitude of the shortest paths for positive and negative tend to balance each other by changing the number of edges or segments to compensate for the changes in the path lengths.

### 3.4 Region Set Enrichment Analysis

RSEA was performed at three levels: whole brain, clustered regions by significance, and clustered regions by connectomics modules. The whole brain analysis (Figure S5) shows FSEC variations across all stages. For EMCI and MCI, most FSEC exhibit increases compared to CN, suggesting a relative higher metabolic demand. For MCI, FSEC (5 for males and 3 for females) decreased compared to CN. For example, decreases in metabolic demand appear in modules encoding for memory and emotions. During AD, there is an increase in the number of FSEC that exhibited decreases, except for motor FSEC, which show an increased metabolic demand relative to CN throughout the disease spectrum. Additionally, auditory senses, memory, and language FSECs display significant were reduced relative to CN for both sexes in AD.

RSEA analysis over regions clustered by significance (Figure S6) revealed a finer level of detail of the changes across the disease spectrum. By performing the analysis in this manner, it is possible to see how regions within each cluster, and how they contribute to SE categories. In EMCI and MCI for both sexes, the changes within each cluster are largely positive, indicating an increase in metabolic demand. In contrast, for LMCI and AD, the alterations are predominantly decreasing within each module, indicating decline in brain FSEC. For example, in the case of memory FSEC, which is associated with cognitive loss in AD, there is an increase within cluster number relative to CN in the early stages. At later stages, this FSEC directionally decreases, which is related to a decline in key brain functions. Lastly, for AD, the FSEC for memory was significant in 7 out of 8 possible categories (75% males and 100% females).

Lastly, RSEA analysis of clustered regions by modules (Figure 3) provides deeper insight into these changes. It shows different alteration levels in each cluster and reveals distinct patterns that ultimately lead to similar outcomes: a decline in auditory senses, cognitive abilities, memory, and language FSEC with an increase in motor FSEC at AD, for both sexes. This suggests that regions follow different trajectories by module and sex, which can be quantified using this method.

**Figure 3.**
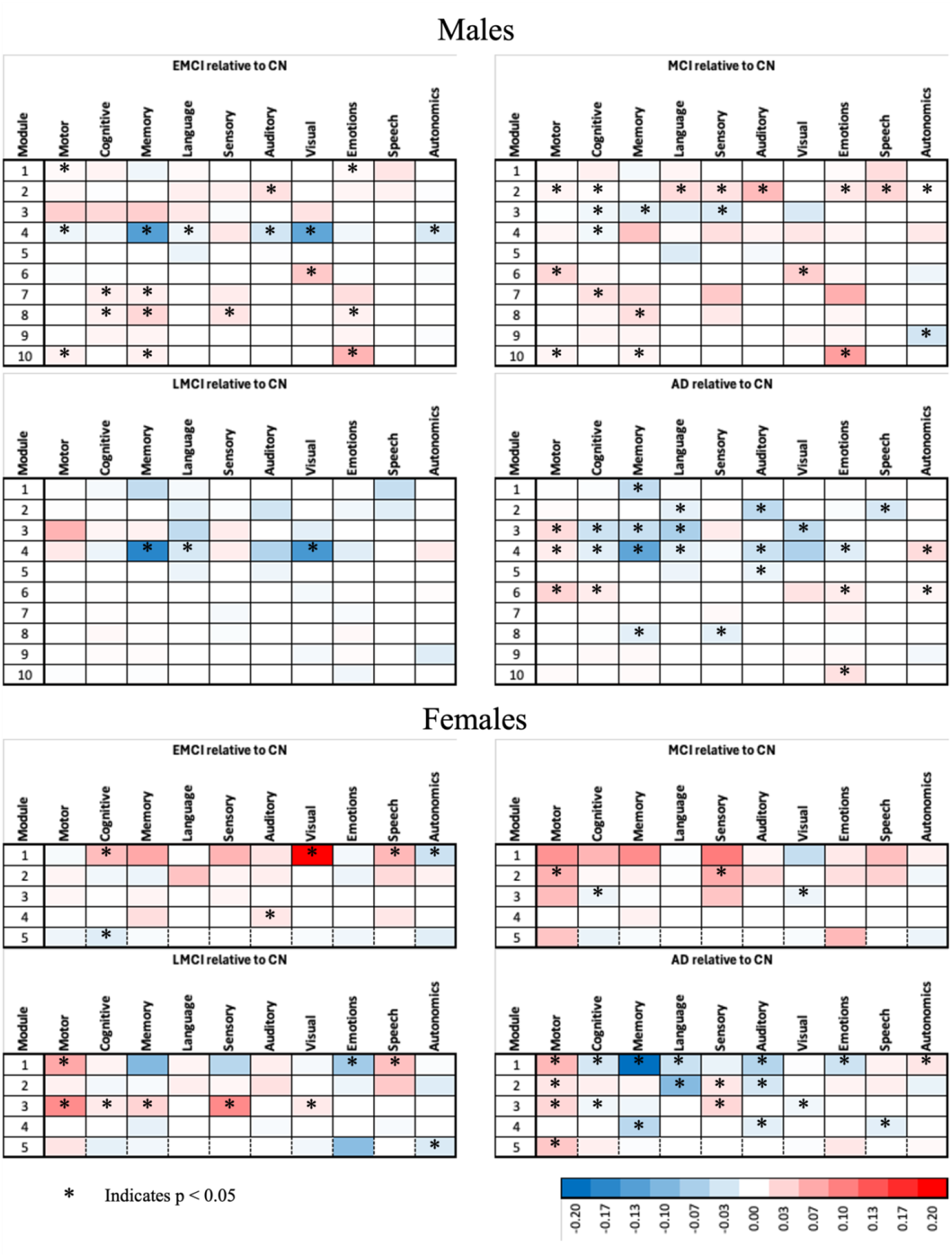
RSEA Performed by Clustering the Regions Using the CN Modules for Each Sex. RSEA performed using the modules obtained from performing the MRCC on the CN cases reveals different alteration levels in each cluster and distinct patterns that ultimately lead to similar outcomes obtained in the previous analysis: a decline in auditory senses, cognitive abilities, memory, and language, accompanied by an increase in motor functions, in both sexes. By performing the analysis by modules, we gain a deeper insight into these changes, enabling a better comprehension of these alterations.

### 3.5 Transcriptomics

Gene expression changes in a system-level framework were identified through WGCNA from the blood transcriptome, with 28 distinct modules of co-expressed genes (Figure S7). To assess functional significance, each of these modules was correlated to the average z-scores of FDG, sex, and disease stage (MCI/AD) of the subjects (Figure 4). We identified two modules significantly associated with MCI (p<0·05)(shown in sienna3, lightgreen), four modules significantly associated with AD (p<0·05)(brown4, red, pink, saddlebrown), and one module significantly associated with FDG (p<0·05)(brown4) (Figure S7). Although grey60 was not significantly associated with FDG or AD and showed only nominal significance with MCI (p<0·1), it exhibited substantial change in the lipid and mitochondrial metabolism biological domains, warranting further investigation. Hence, from all the listed modules, we will focus the analysis on the modules that contain biological domains associated with metabolic changes (MC): red, sienna3, brown4, and grey60 (Figure 4·A).

**Figure 4.**
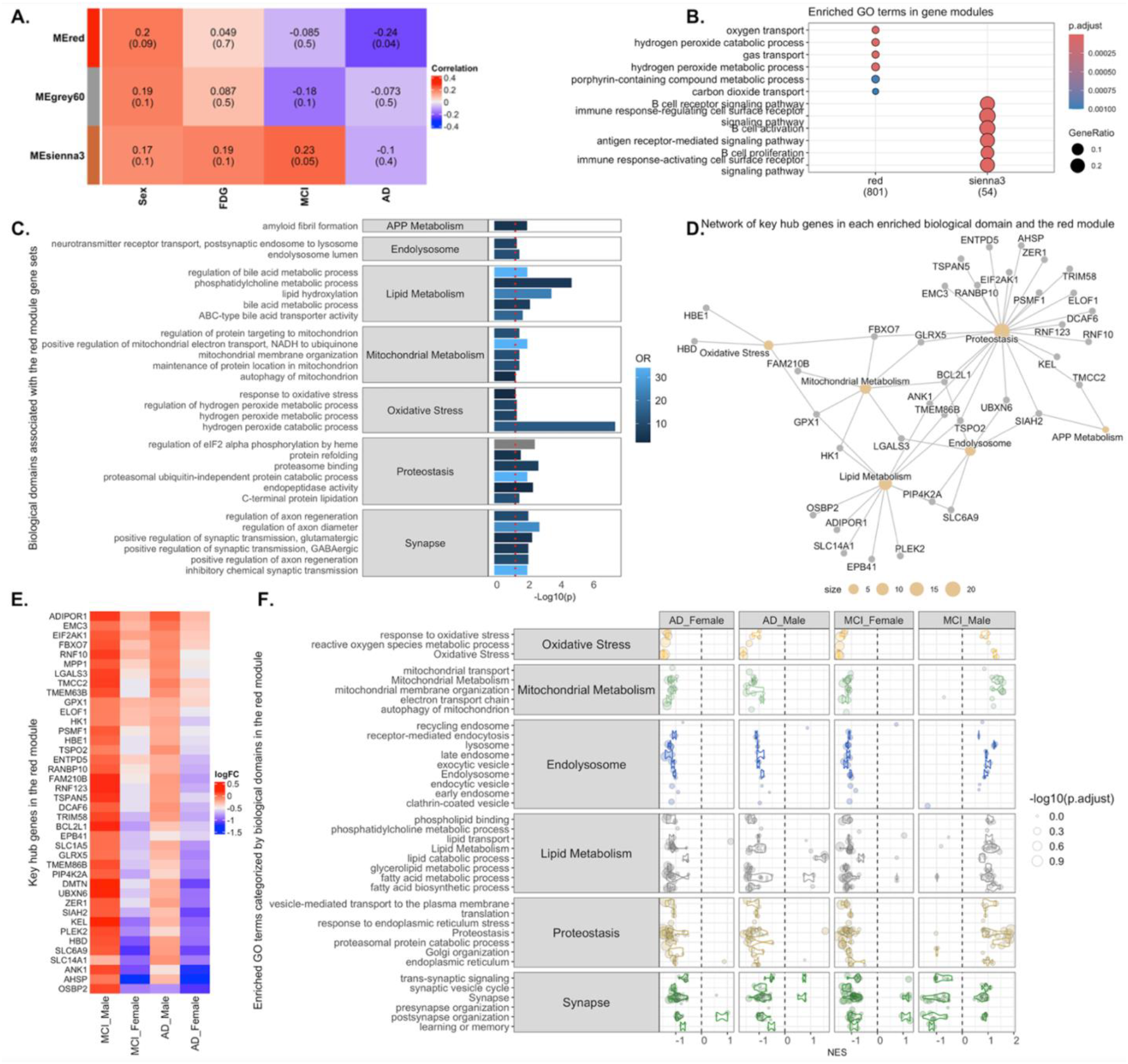
WGCNA Module-trait Associations and Functional Characterization of AD-related Gene Modules. (**A**) Heatmap showing correlations between WGCNA-identified gene modules and traits including average z-scores of FDG uptake, AD status, MCI status, and sex. The red module is significantly associated with AD (p<0·05), the sienna3 module with FDG (p<0·1) and MCI (p<0·05), and the grey60 module with MCI (p=0·1). All three modules show associations with sex (p<0·1). (**B**) Dot plot showing KEGG pathway enrichment analysis results (adjusted p<0·1) for genes in the red and sienna3 modules. (**C**) Bar plot showing enriched AD-related biological domains and resident GO terms in the red module gene set. Each bar represents the strength of enrichment, measured by the odds ratio (OR) and the statistical significance (p-value). (**D**) Network visualization of hub genes within each enriched biological domain in the red module. (**E**) Heatmap of log fold change values for hub genes in MCI and AD cases compared to sex-matched controls. (**F**) Gene set enrichment analysis (GSEA) results for red module genes in MCI male, MCI female, AD male, and AD female subgroups are shown with normalized enrichment scores (NES). Each point is a GO term within the indicated biological domain, the size of the point is scaled by the GSEA adjusted p value, and the normalized enrichment scores (NES) indicates if the GO term is up- or downregulated.

**Figure 5.**
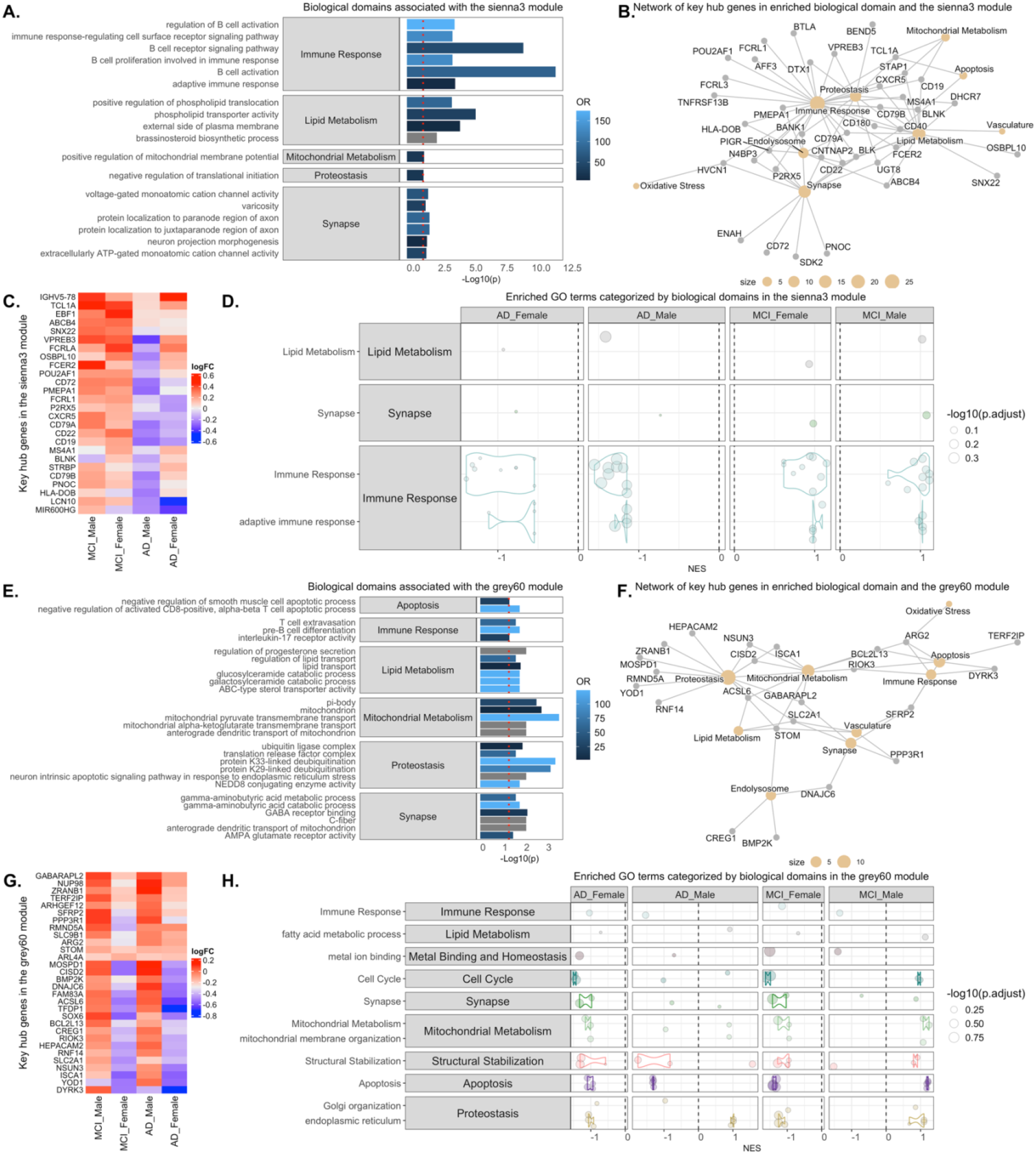
Functional Characterization of the FDG- and MCI-associated WGCNA Modules. Bar plot showing enriched AD-related biological domains and resident GO terms in **(A)** the sienna3 module and (**E**) the grey60 gene set. Each bar represents the strength of enrichment, measured by the odds ratio (OR) and the statistical significance (p-value). Network visualization of hub genes within each enriched biological domain in **(B)** the sienna3 and **(F)** the grey60 module. Heatmap of log fold change values for (**C**) sienna3 and (**G**) grey60 module hub genes in MCI and AD cases compared to sex-matched controls. GSEA results for **(D)** sienna3 module genes and **(H)** grey60 module genes in MCI male, MCI female, AD male, and AD female subgroups. Each point is a GO term within the indicated biological domain, the size of the point is scaled by the GSEA adjusted p-value, and the normalized enrichment scores (NES) indicates if the GO term is up- or downregulated.

Enriched GO terms for the red module identified enrichment in oxygen, carbon dioxide, and gas transport, as well as hydrogen peroxide and porphyrin-containing compound metabolic processes. On the other hand, the sienna3 module identified enrichment in B cell receptor, antigen receptor-mediated, and immune response-activating cell surface receptor signaling pathways, B cell proliferation, and B cell activation (Figure 4·B).

Next, we evaluated the enrichment of biological domains within MC modules (Figures 4·C-F/5/S8) and found that GO-terms associated with lipid and mitochondrial metabolism, oxidative stress, proteostasis, and synapse biological domains were significantly enriched (p<0·05). By examining the biological domains of each module, we highlight the following changes: mitochondrial metabolism and membrane organization (red, grey60), lysosome (brown4), lipid metabolism (red, sienna3), fatty acid metabolic process (red, grey60), Golgi organization (red, grey60, brown4), endoplasmic reticulum (red, grey60, brown4), synapse (red, sienna3, grey60, brown4), immune response (sienna3, grey60, brown4), metal ion binding and structural stabilization (grey60, brown4), and cytokines (brown4). For a complete list of changes, see Figures 4·C-F/5/S8. These biological domains are heavily related to cellular homeostasis, cellular regulating processes, cellular signaling, and cellular synthesis, with dysregulation linked to neurodegenerative disorders. For example, cytokines dysregulation is linked with chronic inflammation, autoimmune disorders, and impaired immune response; endoplasmic reticulum dysregulation is linked with metabolic disorders, increased oxidative stress, and disruption of lipid metabolism.

### 3.6 Clinical Cognitive Assessment

Table S7 presents the results of Pearson’s correlation between the connectomics metrics and CCA scores. When comparing these measures, positive nodal strength was identified as the connectomics metric with significant correlations relative to various task scores of each CCA. The relationship was positively correlated within the range of 0·5862 to 0·8779. Other connectomics metrics such as density, degrees, clustering, and negative nodal strengths also exhibited positive correlations with CCA scores (Table S7). In contrast, the number of path segments and path lengths displayed a tendency towards negative correlations relative to CCA scores.

## 4. DISCUSSION

By analyzing the network-level metabolic metrics, it was observed that their values an oscillation between AD stages. For instance, network density and degree distribution for both sexes follow a ‘W’ pattern, where there is a decrease in values followed by an increase. This indicates that the brain undergoes restructuring as the disease progresses since these metrics measure the number of connections and how these nodes are connected, which is consistent with brain metabolic reprogramming theory^4^, and suggests changes in the efficiency of communication between regions. Additionally, evaluating the clustering coefficient and nodal strength shows that nodes become more decentralized as the disease progresses (Figure 1·A-E), suggesting alterations in the energy dependent connectivity networks related to disease progression.

Similarly, an assessment of changes in the WTN reveals that these networks become more sparse as the disease progresses (Figure 1·F-O). Consistent with these changes, analysis of MRCC modules indicates that the number and region composition within each module vary across the disease spectrum. Notably, males and females exhibit different patterns: in males, the number of modules decreases (indicating network collapse); whereas in females, the number of modules increases (indicating a more fractured network), suggesting sexual dimorphism in metabolic brain module with disease state. Importantly, these changes are consistent with our previous reports in mouse models of AD,^9^ illustrating the universality of this approach. Collectively, these findings suggest that brain regions prune network paths as the disease advances, which is consistent with the idea that the brain undergoes a restructuring process in an attempt to maintain normal functions.^4,12,13,18^

In the current study, positive network changes represent excitatory activity of neuronal firing, while negative network activity represents inhibitor activity of neuronal firing, effectively dampening neural activity and regulating excitatory activity to prevent disorders like epilepsy or anxiety.^40^ Importantly, regulating Excitation/Inhibition (E/I) balance in the brain is crucial for maintaining proper neural function, regulating information processing, and preventing excessive or inadequate neural activity.^41^ In this context, positive and negative covariance paths in metabolic covariance matrices represent excitatory and inhibitory signals, respectively, since positive entries represent synchronized positive correlations between brain regions, while negative entries indicate an inverse relationship, aligning with inhibitory signals. Under these conditions, positive and negative paths should be minimized to avoid an E/I imbalance that leads to pathophysiological states.^40,42^

We also observe variations in the number of paths, path lengths, and edge counts across the ADRD spectrum; however, the difference between the ratios of positive and negative path lengths and their corresponding number of segments remains constant. This observation aligns with the brain energy failure^21^ and reorganization^4,5^ theories, which states that under bioenergetic stress, the brain will attempt to support energy demands by performing fuel switching; however, under extreme loads the brain will be forced to reorganize brain networks to continue to support normal function.^16,17^ By comparing the distances between pairwise regions, reveals a decrease in distances during MCI, consistent with the aforementioned theories. This decrease in path length implies an increase in efficiency in the network, which is associated with an increased metabolic activity (hypermetabolism) in the brain, while a decrease in path length is associated with a loss of efficiency as the brain begins to fail (hypometabolism). The current connectomics and path analysis suggests that in the transition from EMCI to MCI the brain increases its metabolic demand in support of increasing pathophysiology. By contrast, during the transition from MCI to LMCI is when energetic demand exceeds the functional reserve, leading to network failure, and brain metabolic instability.^6,7,43^ Importantly, these network behaviors align with previous reports, which describe the metabolic alterations in the critical brain regime hypothesis, which proposes that optimal brain function occurs near a phase transition (an intermediate stage where neuronal networks shift between different activity states, enhancing information processing and adaptability);^42^ and an extended critical regime phase (also referred as Griffith’s Phase; a phase where neuronal networks exhibit critical-like properties for an extended period of time)^44^ is associated with AD due to excessive excitation correlated to hyper-synchronization and cognitive decline.^13,44,45^

To explore this in the context of brain function, RSEA permits quantification of changes in FSEC relative to a reference group. This novel method allows us to examine how metabolic alterations between each phase impact various FSEC. Across all levels of whole brain analysis, motor, memory, and language FSECs were significant across the ADRD spectrum (Figures 3/S5/S6). In an effort to provide additional refinement of the FSEC changes through ADRD progression, RSEAs analysis was applied to MRCC modules. For instance, significant cognitive FSEC changes were observed at the module level across the disease spectrum (Figure 3), but they are not discernible at the whole brain level (Figure S5). Moreover, RSEA matrices predominantly exhibit increases relative to CN in the early stages, but this trend was reversed during the later stages of the disease, and align with clinical variations in metabolism and cognitive decline described in the AD population.^12,13,19,21^

Transcriptomics revealed changes that support our findings by identifying alterations in lipid and mitochondrial metabolism, oxidative stress, proteostasis, and synapse (Figure 4/5/S8). Also, in a more granular manner, it revealed changes in processes at a cellular level that are linked to neurodegenerative disorders. For example, lipid, mitochondrial, and fatty acid metabolism changes may indicate a shift in the metabolic processes within cells, often reflecting a move from normal metabolic homeostasis toward alternative pathways that are less efficient or more damaging to cellular health. Lysosome alterations may lead to amyloid-β accumulation due to impaired clearance, mitochondrial dysfunction, increased oxidative stress, and chronic inflammation, which aligns with the metabolic reprogramming theory.^4,5^ Dysregulation in Golgi organization and structural stabilization may suggest impairment of neuronal physiology due to the deficits in protein processing, stress signaling, and microtubule and vesicular transport impairment; hence, it might be an independent process that contributes to disease progression and not just a byproduct of cell death. Cytokines are signaling proteins secreted by immune cells (i.e., microglia, macrophages) that are important in mediating and regulating immune and inflammatory responses and maintaining central nervous system homeostasis. Hence, cytokine dysregulation activates microglia, which release pro-inflammatory cytokines (i.e., TNF-α, IL-1β, IL-6) as an initial/acute protective response, but chronic activation leads to chronic neuroinflammation, which is a central mechanism underlying neurodegenerative progression. Lastly, endoplasmic reticulum (ER) serves a role in protein folding, calcium homeostasis, and quality control within neuros; however, when the ER is under stress or dysfunctional like in AD, it can lead to accumulation of misfolded proteins (i.e., amyloid-β, hyperphosphorylated tau tangles) which overwhelms the ER’s capacity and ends contributing to the onset and progression of AD. These findings align with the energy failure^21^ and metabolic reprogramming theories.^4,5^

To relate our connectomics and shortest path findings with CCA, we performed covariate analysis, which revealed that the current analysis retains the ability to assess disease changes which may be beyond the ability of CCA tests to detect disease early. This disconnect could be attributed to the fact that the behavior across the disease spectrum exhibits a non-linear relationship with CCA response, which is a limitation of CCA scores. Hence, connectomics and shortest path analysis may provide a complementary analysis to CCA.

One important limitation of this study is that the analysis is conducted at a macro-level, as we are analyzing unilateral regional data, which may potentially overlook bilateral changes and reveal more profound changes. Additionally, these limitations may affect the shortest path length analysis, because these covariance matrices do not directly represent synaptic connection, but rather describe statistical interdependencies. Furthermore, this study was conducted using a single imaging mode, and to encompass the entire disease spectrum, multimodal imaging should be considered, which highlights the need for integrated multimodal brain connectomics modeling approach.

In conclusion, this study demonstrated the value of analyzing metabolic alterations using connectomics and path length metrics. The results from these metrics support the idea that the brain undergoes a network reorganization as a compensatory mechanism of pathological disruption. Moreover, path lengths between brain regions change as AD advances, decreasing in MCI but increasing in AD. This suggests a decrease in the efficiency of the connectivity network, which may contribute to cognitive decline. RSEA analysis revealed significant functional changes in motor, memory, language, and cognition, highlighting the impact of these metabolic variations on the subject’s FSEC. Integrating multimodal data will be critical for developing a more comprehensive framework that captures metabolic, functional, and structural changes, thereby providing deeper insights into disease progression and potential therapeutic targets.

## ACKNOWLEDGEMENTS

Data collection and sharing for this project was funded by the Alzheimer’s Disease Neuroimaging Initiative (ADNI) (National Institutes of Health Grant U01 AG024904) and DOD ADNI (Department of Defense award number W81XWH-12-2-0012). ADNI is funded by the National Institute on Aging, the National Institute of Biomedical Imaging and Bioengineering, and through generous contributions from the following: AbbVie, Alzheimer’s Association; Alzheimer’s Drug Discovery Foundation; Araclon Biotech; BioClinica, Inc.; Biogen; Bristol-Myers Squibb Company; CereSpir, Inc.; Cogstate; Eisai Inc.; Elan Pharmaceuticals, Inc.; Eli Lilly and Company; EuroImmun; F. Hoffmann-La Roche Ltd and its affiliated company Genentech, Inc.; Fujirebio; GE Healthcare; IXICO Ltd.; Janssen Alzheimer Immunotherapy Research & Development, LLC.; Johnson & Johnson Pharmaceutical Research & Development LLC.; Lumosity; Lundbeck; Merck & Co., Inc.; Meso Scale Diagnostics, LLC.; NeuroRx Research; Neurotrack Technologies; Novartis Pharmaceuticals Corporation; Pfizer Inc.; Piramal Imaging; Servier; Takeda Pharmaceutical Company; and Transition Therapeutics. The Canadian Institutes of Health Research is providing funds to support ADNI clinical sites in Canada. Private sector contributions are facilitated by the Foundation for the National Institutes of Health (www.fnih.org). The grantee organization is the Northern California Institute for Research and Education, and the study is coordinated by the Alzheimer’s Therapeutic Research Institute at the University of Southern California. ADNI data are disseminated by the Laboratory for Neuro Imaging at the University of Southern California. We thank Dorsey Faught from MIM Software Inc who helped with the use and development of workflows in MIM.

## AUTHOR’S CONTRIBUTIONS

JAKCC contributed to the study design, data collection and management, data analysis and interpretation, data quality assessment, literature review, and manuscript drafting. SAC participated in study design, data collection and management, data analysis, and quality assessment. RSP provided transcriptomics analysis and interpretation, and helped write the manuscript. ORS did literature search for functional analysis of the brain regions. PS provided statistical and image analysis advice, participated in data interpretation, and helped write the manuscript. PRT contributed to the study design, data analysis and interpretation, and helped write the manuscript. JAKCC, PRT, and SAC had access to all raw data. RSP had access to raw genetic data. JAKCC, PRT, and SAC verified the data and results. All authors helped revise the manuscript. All authors had final responsibility for the decision to submit for publication.

## DATA SHARING

The data that support the findings of this study were obtained from the Alzheimer’s Disease Neuroimaging Initiative (ADNI), which is available from the ADNI database (https://adni.loni.usc.edu) upon registration and compliance with the data use agreement.

## ETHICS COMMITTEE APPROVAL

Ethics approval was not required for this study.

## REFERENCES

1. 2024 Alzheimer’s disease facts and figures. Alzheimers Dement. 2024;20(5):3708–821.

2. Nandi A, Counts N, Broker J, Malik S, Chen S, Han R, et al. Cost of care for Alzheimer’s disease and related dementias in the United States: 2016 to 2060. NPJ Aging. 2024;10(1):13.

3. Karch CM, Cruchaga C, Goate AM. Alzheimer’s disease genetics: from the bench to the clinic. Neuron. 2014;83(1):11–26.

4. Demetrius LA, Eckert A, Grimm A. Sex differences in Alzheimer’s disease: metabolic reprogramming and therapeutic intervention. Trends Endocrinol Metab. 2021;32(12):963–79.

5. Demetrius LA, Magistretti PJ, Pellerin L. Alzheimer’s disease: the amyloid hypothesis and the Inverse Warburg effect. Front Physiol. 2014;5:522.

6. Onos KD, Lin PB, Pandey RS, Persohn SA, Burton CP, Miner EW, et al. Assessment of neurovascular uncoupling: APOE status is a key driver of early metabolic and vascular dysfunction. Alzheimers Dement. 2024;20(7):4951–69.

7. Chong Chie JAK, Persohn SA, Pandey RS, Simcox OR, Salama P, Territo PR, et al. Neuro-Metabolic and Vascular Dysfunction as an Early Diagnostic for Alzheimer’s Disease and Related Dementias. bioRxiv. 2025.

8. Nordberg A, Rinne JO, Kadir A, Langstrom B. The use of PET in Alzheimer disease. Nat Rev Neurol. 2010;6(2):78–87.

9. Chumin EJ, Burton CP, Silvola R, Miner EW, Persohn SC, Veronese M, et al. Brain metabolic network covariance and aging in a mouse model of Alzheimer’s disease. Alzheimers Dement. 2024;20(3):1538–49.

10. Cohen AD, Klunk WE. Early detection of Alzheimer’s disease using PiB and FDG PET. Neurobiol Dis. 2014;72 Pt A:117–22.

11. Macdonald IR, DeBay DR, Reid GA, O’Leary TP, Jollymore CT, Mawko G, et al. Early detection of cerebral glucose uptake changes in the 5XFAD mouse. Curr Alzheimer Res. 2014;11(5):450–60.

12. Johnson SC, Christian BT, Okonkwo OC, Oh JM, Harding S, Xu G, et al. Amyloid burden and neural function in people at risk for Alzheimer’s Disease. Neurobiol Aging. 2014;35(3):576–84.

13. Yi D, Lee DY, Sohn BK, Choe YM, Seo EH, Byun MS, et al. Beta-amyloid associated differential effects of APOE epsilon4 on brain metabolism in cognitively normal elderly. Am J Geriatr Psychiatry. 2014;22(10):961–70.

14. Perovnik M, Tang CC, Namias M, Eidelberg D, Alzheimer’s Disease Neuroimaging I. Longitudinal changes in metabolic network activity in early Alzheimer’s disease. Alzheimers Dement. 2023;19(9):4061–72.

15. Veronese M, Moro L, Arcolin M, Dipasquale O, Rizzo G, Expert P, et al. Covariance statistics and network analysis of brain PET imaging studies. Sci Rep. 2019;9(1):2496.

16. Fornito A, Zalesky A, Bullmore E. Fundamentals of brain network analysis: Academic press; 2016.

17. Bassett DS, Sporns O. Network neuroscience. Nat Neurosci. 2017;20(3):353–64.

18. Sala A, Lizarraga A, Caminiti SP, Calhoun VD, Eickhoff SB, Habeck C, et al. Brain connectomics: time for a molecular imaging perspective? Trends Cogn Sci. 2023;27(4):353–66.

19. Titov D, Diehl-Schmid J, Shi K, Perneczky R, Zou N, Grimmer T, et al. Metabolic connectivity for differential diagnosis of dementing disorders. J Cereb Blood Flow Metab. 2017;37(1):252–62.

20. Perneczky R, Drzezga A, Diehl-Schmid J, Li Y, Kurz A. Gender differences in brain reserve : an (18)F-FDG PET study in Alzheimer’s disease. J Neurol. 2007;254(10):1395–400.

21. Zhang X, Alshakhshir N, Zhao L. Glycolytic Metabolism, Brain Resilience, and Alzheimer’s Disease. Front Neurosci. 2021;15:662242.

22. Rodgers ZB, Detre JA, Wehrli FW. MRI-based methods for quantification of the cerebral metabolic rate of oxygen. J Cereb Blood Flow Metab. 2016;36(7):1165–85.

23. Iwatsubo T. [Alzheimer’s disease Neuroimaging Initiative (ADNI)]. Nihon Rinsho. 2011;69 Suppl 8:570–4.

24. Chong Chie JAK, Persohn SC, Miner EW, Burton CP, Salama P, Territo PR, editors. Total Variation Based 2D Image Registration of Post-Mortem Mouse Brain Images. 2024 IEEE International Symposium on Biomedical Imaging (ISBI); 2024: IEEE.

25. Cavusoglu B, Durak H. The Effect of Patient Age on Standardized, Uptake Value-Hounsfield Unit Values of Male Genitourinery Structures In F-18 FDG PET/CT. Mol Imaging Radionucl Ther. 2011;20(3):104–7.

26. Kinahan PE, Fletcher JW. Positron emission tomography-computed tomography standardized uptake values in clinical practice and assessing response to therapy. Semin Ultrasound CT MR. 2010;31(6):496–505.

27. Nugent S, Croteau E, Potvin O, Castellano CA, Dieumegarde L, Cunnane SC, et al. Selection of the optimal intensity normalization region for FDG-PET studies of normal aging and Alzheimer’s disease. Sci Rep. 2020;10(1):9261.

28. Toledo JB, Bjerke M, Da X, Landau SM, Foster NL, Jagust W, et al. Nonlinear Association Between Cerebrospinal Fluid and Florbetapir F-18 beta-Amyloid Measures Across the Spectrum of Alzheimer Disease. JAMA Neurol. 2015;72(5):571–81.

29. Rubinov M, Sporns O. Complex network measures of brain connectivity: uses and interpretations. Neuroimage. 2010;52(3):1059–69.

30. Mucha PJ, Richardson T, Macon K, Porter MA, Onnela JP. Community structure in time-dependent, multiscale, and multiplex networks. Science. 2010;328(5980):876–8.

31. Jeub LGS, Sporns O, Fortunato S. Multiresolution Consensus Clustering in Networks. Sci Rep. 2018;8(1):3259.

32. Costantini G, Perugini M. Generalization of clustering coefficients to signed correlation networks. PLoS One. 2014;9(2):e88669.

33. Turner MH, Mann K, Clandinin TR. The connectome predicts resting-state functional connectivity across the Drosophila brain. Curr Biol. 2021;31(11):2386–94 e3.

34. Seguin C, van den Heuvel MP, Zalesky A. Navigation of brain networks. Proc Natl Acad Sci U S A. 2018;115(24):6297–302.

35. Bordbar A, Nagarajan H, Lewis NE, Latif H, Ebrahim A, Federowicz S, et al. Minimal metabolic pathway structure is consistent with associated biomolecular interactions. Mol Syst Biol. 2014;10(7):737.

36. Langfelder P, Horvath S. WGCNA: an R package for weighted correlation network analysis. BMC Bioinformatics. 2008;9:559.

37. Yu G, Wang LG, Han Y, He QY. clusterProfiler: an R package for comparing biological themes among gene clusters. OMICS. 2012;16(5):284–7.

38. Cary GA, Wiley JC, Gockley J, Keegan S, Amirtha Ganesh SS, Heath L, et al. Genetic and multi-omic risk assessment of Alzheimer’s disease implicates core associated biological domains. Alzheimers Dement (N Y). 2024;10(2):e12461.

39. Subramanian A, Tamayo P, Mootha VK, Mukherjee S, Ebert BL, Gillette MA, et al. Gene set enrichment analysis: a knowledge-based approach for interpreting genome-wide expression profiles. Proc Natl Acad Sci U S A. 2005;102(43):15545–50.

40. Sohal VS, Rubenstein JLR. Excitation-inhibition balance as a framework for investigating mechanisms in neuropsychiatric disorders. Mol Psychiatry. 2019;24(9):1248–57.

41. Zhou S, Yu Y. Synaptic E-I Balance Underlies Efficient Neural Coding. Front Neurosci. 2018;12:46.

42. Javed E, Suarez-Mendez I, Susi G, Roman JV, Palva JM, Maestu F, et al. A Shift Toward Supercritical Brain Dynamics Predicts Alzheimer’s Disease Progression. J Neurosci. 2025;45(9).

43. Kotredes KP, Pandey RS, Persohn S, Elderidge K, Burton CP, Miner EW, et al. Characterizing molecular and synaptic signatures in mouse models of late-onset Alzheimer’s disease independent of amyloid and tau pathology. Alzheimers Dement. 2024;20(6):4126–46.

44. Moretti P, Munoz MA. Griffiths phases and the stretching of criticality in brain networks. Nat Commun. 2013;4:2521.

45. Buendia V, Villegas P, Burioni R, Munoz MA. The broad edge of synchronization: Griffiths effects and collective phenomena in brain networks. Philos Trans A Math Phys Eng Sci. 2022;380(2227):20200424.

