## Supplemental Figures S1-S8 for "Multiscale Metabolic Covariance Networks Uncover Stage-Specific Biomarker Signatures Across the Alzheimer’s Disease Continuum"

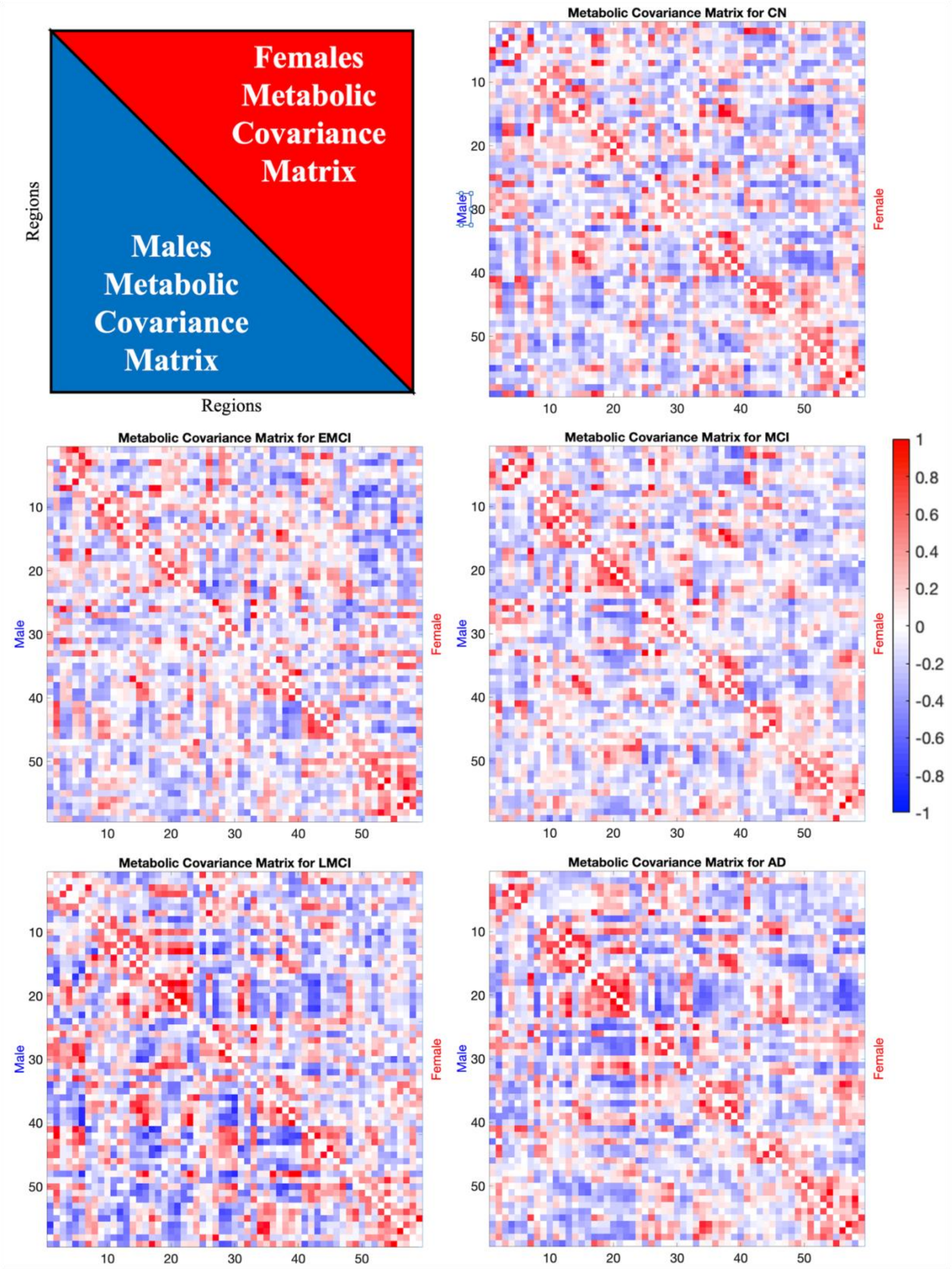


**Supplemental Figure S1 – Unthresholded Metabolic Covariance Matrices.**
(**A**) General overlay of the matrices where the lower triangular matrix and upper triangular matrix are the metabolic covariance matrices for males and females, respectively. The results for each disease stage are shown in: (**B**) CN, (**C**) EMCI, (**D**) MCI, (**E**) LMCI, and (**F**) AD. Covariances were calculated using Pearson correlation of regional SUVR values across each subject within each group. Thresholded matrices for $p<0{\cdot}05$ are shown in Figure S2. Complete names of the regions are found in Table S3.

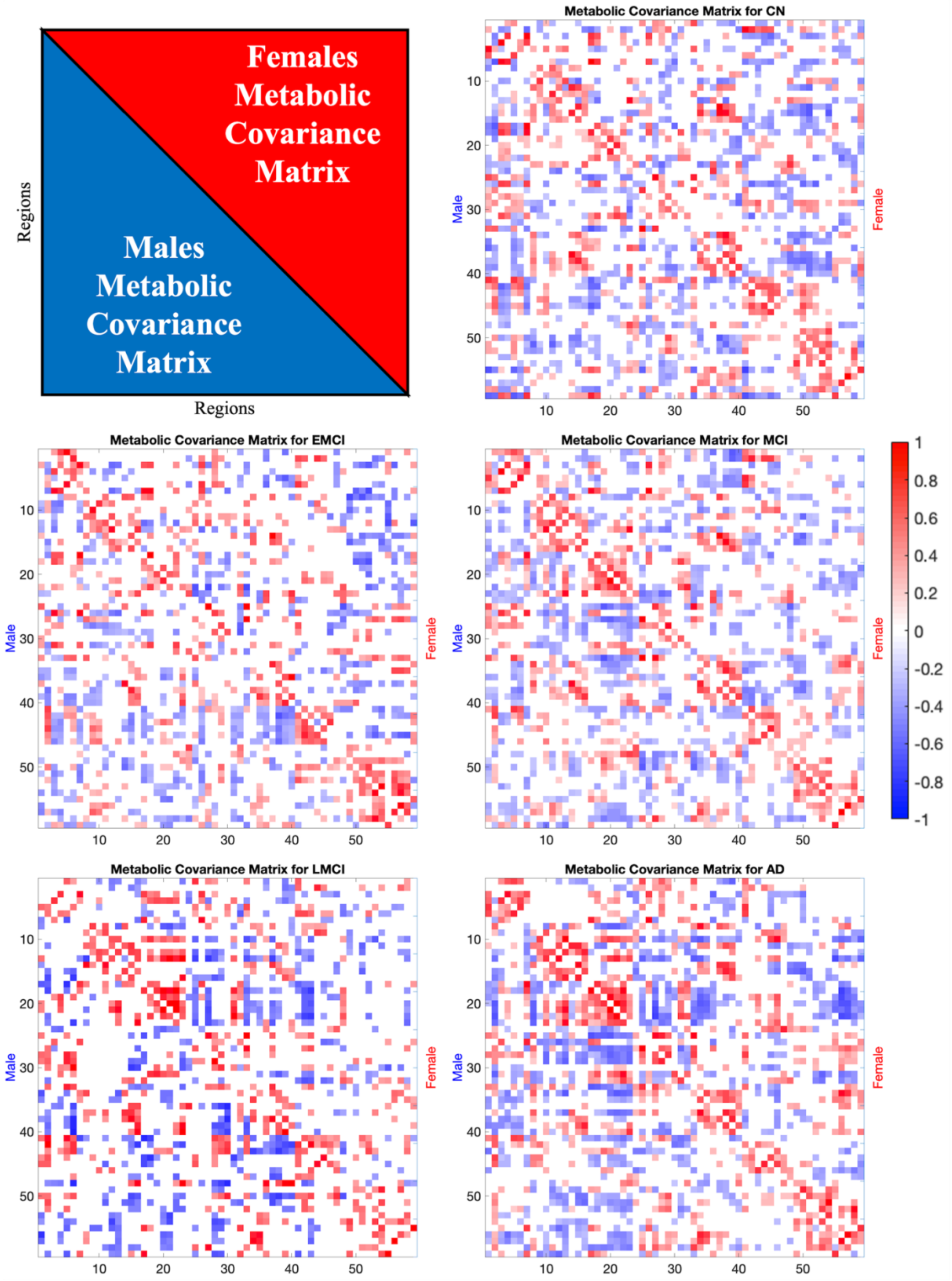


**Supplemental Figure S2 – Metabolic Thresholded Connectomics Matrices p<0·05.**
(**A**) General Overlay, (**B**) CN, (**C**) EMCI, (**D**) MCI, (**E**) LMCI, and (**F**) AD. The lower triangular matrix and upper triangular matrix are the metabolic covariance matrices for males and females, respectively. Complete names of the regions are found in Table S3. As the disease progresses, the matrices become sparse and fragmented, especially in females.

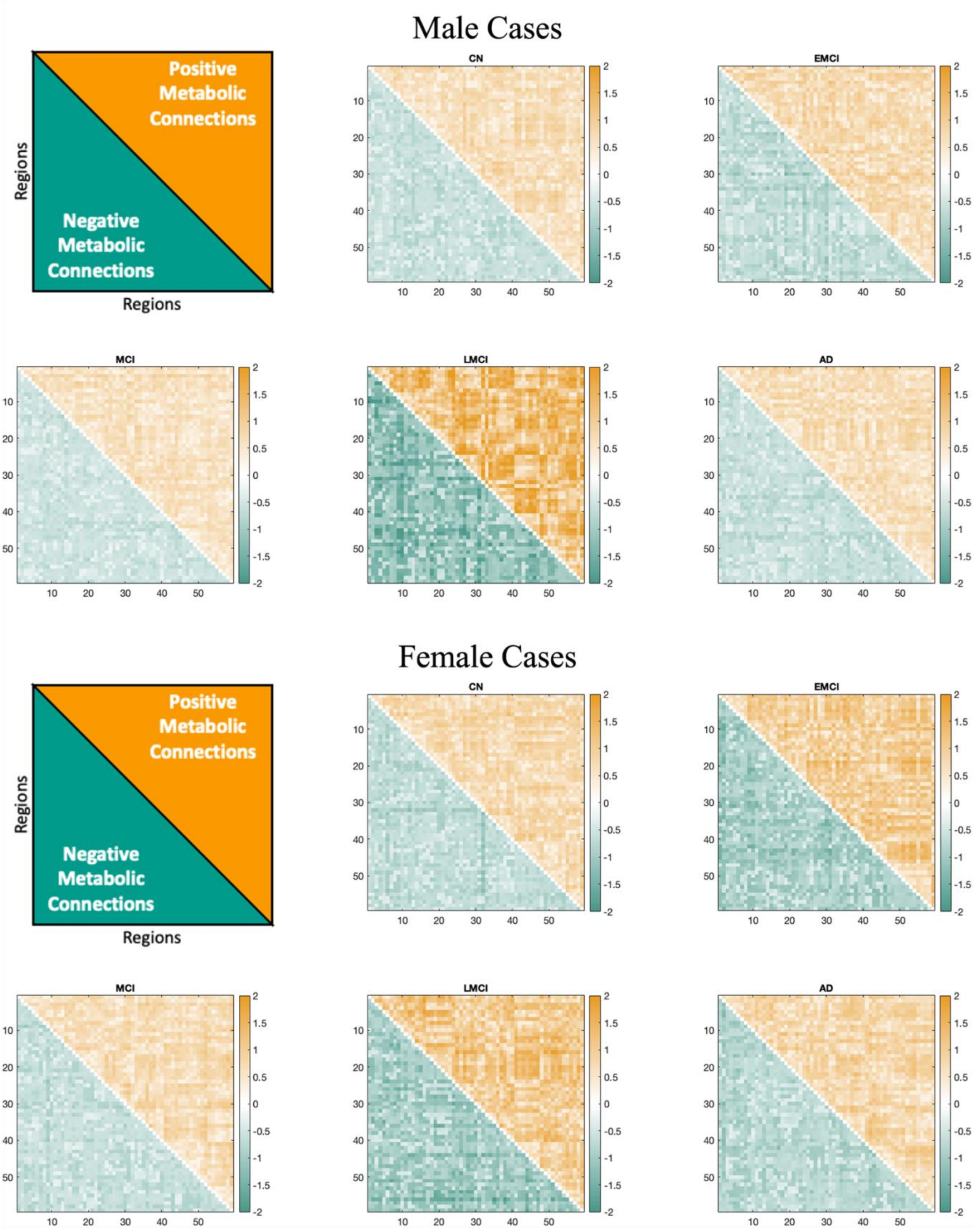


**Supplemental Figure S3 – Shortest Path Lengths for Positive and Negative Pathways.**
Each entry of the distance matrices represents the shortest path length between regions for positive connections in the upper triangular matrix and negative connections in the lower triangular matrix. Diagonal values represent self-connections and are set to 0. Qualitative analysis shows the largest distance length increase during the LMCI phase for both sexes.

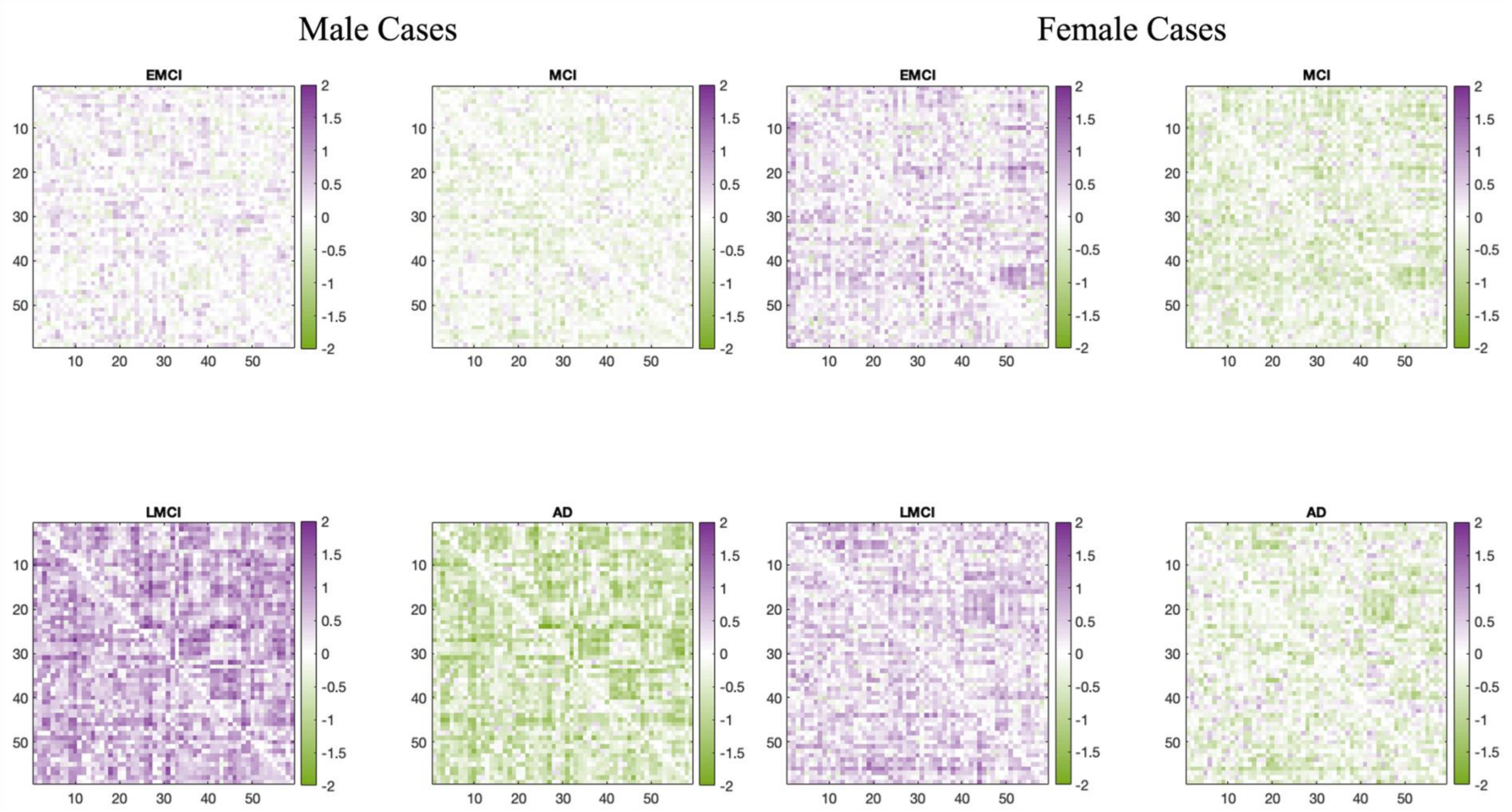


**Supplemental Figure S4 – Comparisons of the Pathways Relative to the Previous Stage.**
In comparison to the preceding disease stage, EMCI and LMCI exhibit increased path lengths compared to their respective prior stages. Conversely, MCI and AD demonstrate decreased path lengths. Similar to the comparison relative to CN, the variations in the path lengths suggest alterations in the energetic balance.

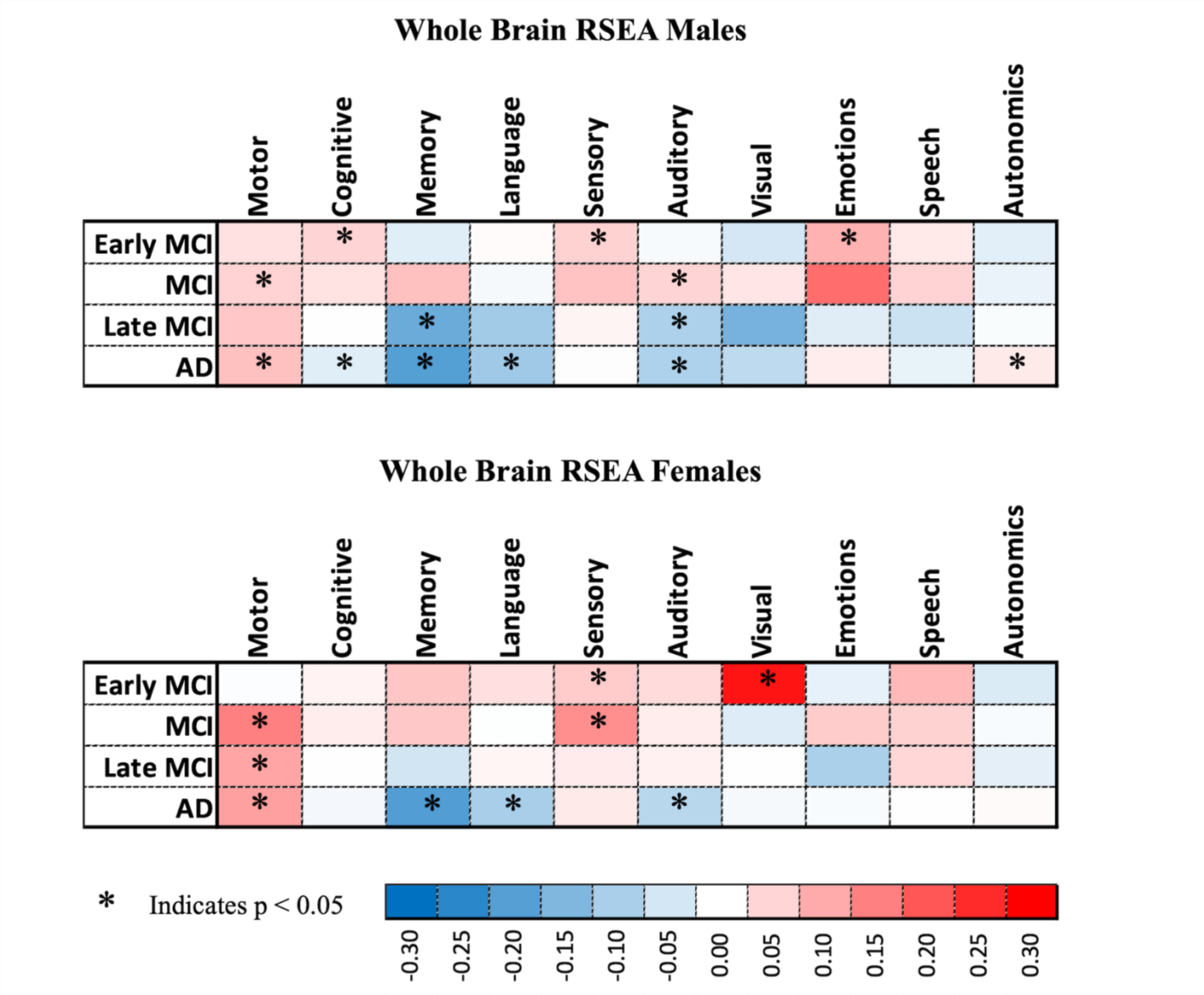


**Supplemental Figure S5 – Whole Brain RSEAs.**

RSEA reveals that the brain's metabolic demand increases during the early stages and decreases during the late stages. Both sexes experience significant changes in language, memory, and motor functions across the disease spectrum. Notably, while cognition shows no significant changes in females, this could be attributed to the limitations of the current analysis method over the whole brain; suggesting that a more granular analysis is required.

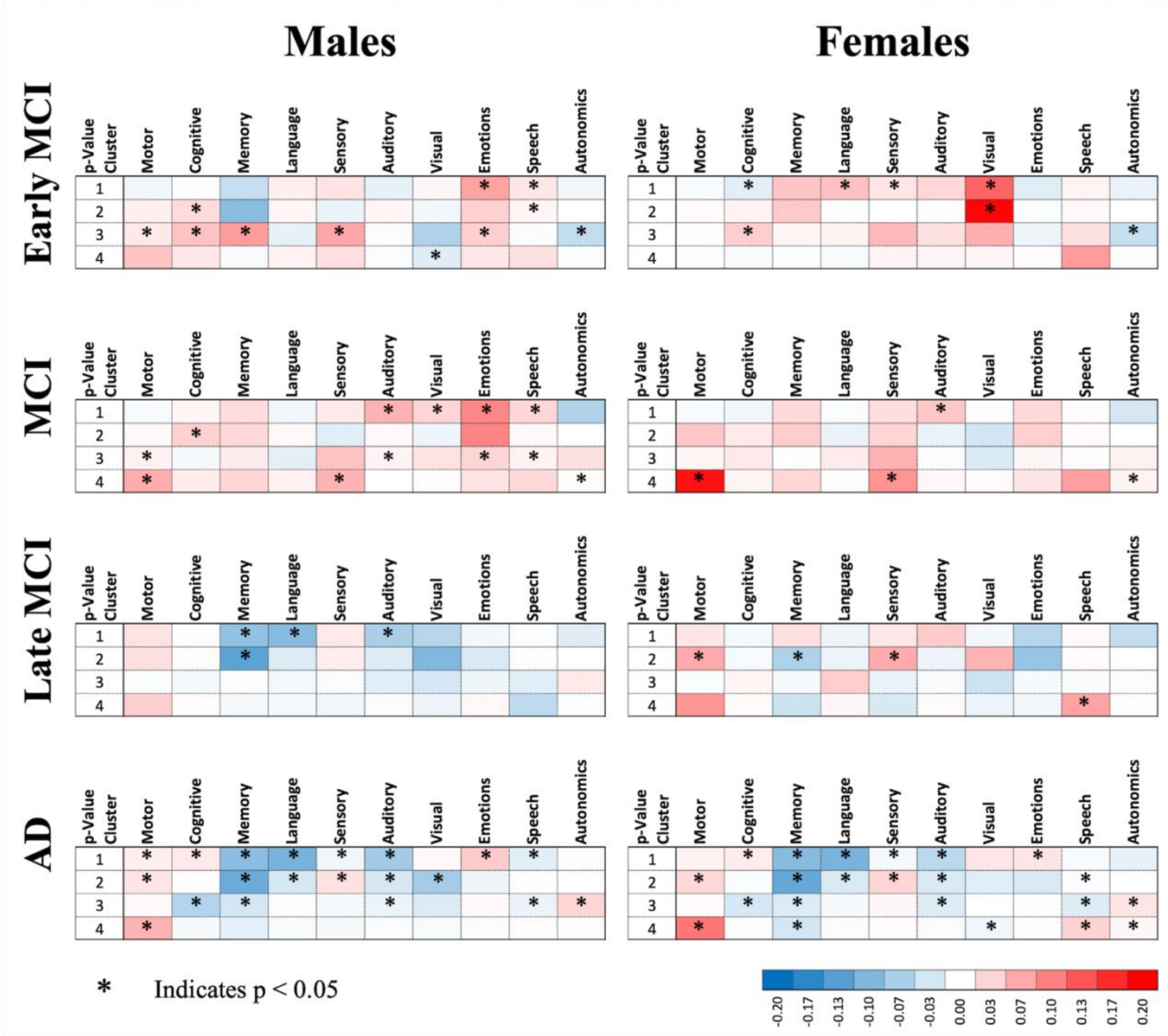


**Supplemental Figure S6 – RSEA Performed by Clustering Regions by Significance Changes.**
Significance was determined using Student's t-test (p<0·05). Subsequently, the p-values were sorted in ascending order, and brain regions were divided into four clusters based on this order: cluster 1 comprises regions 1 through 15, cluster 2 includes regions 16 to 30, cluster 3 contains regions 31 to 45, and cluster 5 comprises regions 46 to 59. RSEA analysis reveals a similar pattern to the whole brain RSEA analysis. However, in addition to the previously identified functional domains, cognition shows as a significant domain in AD.

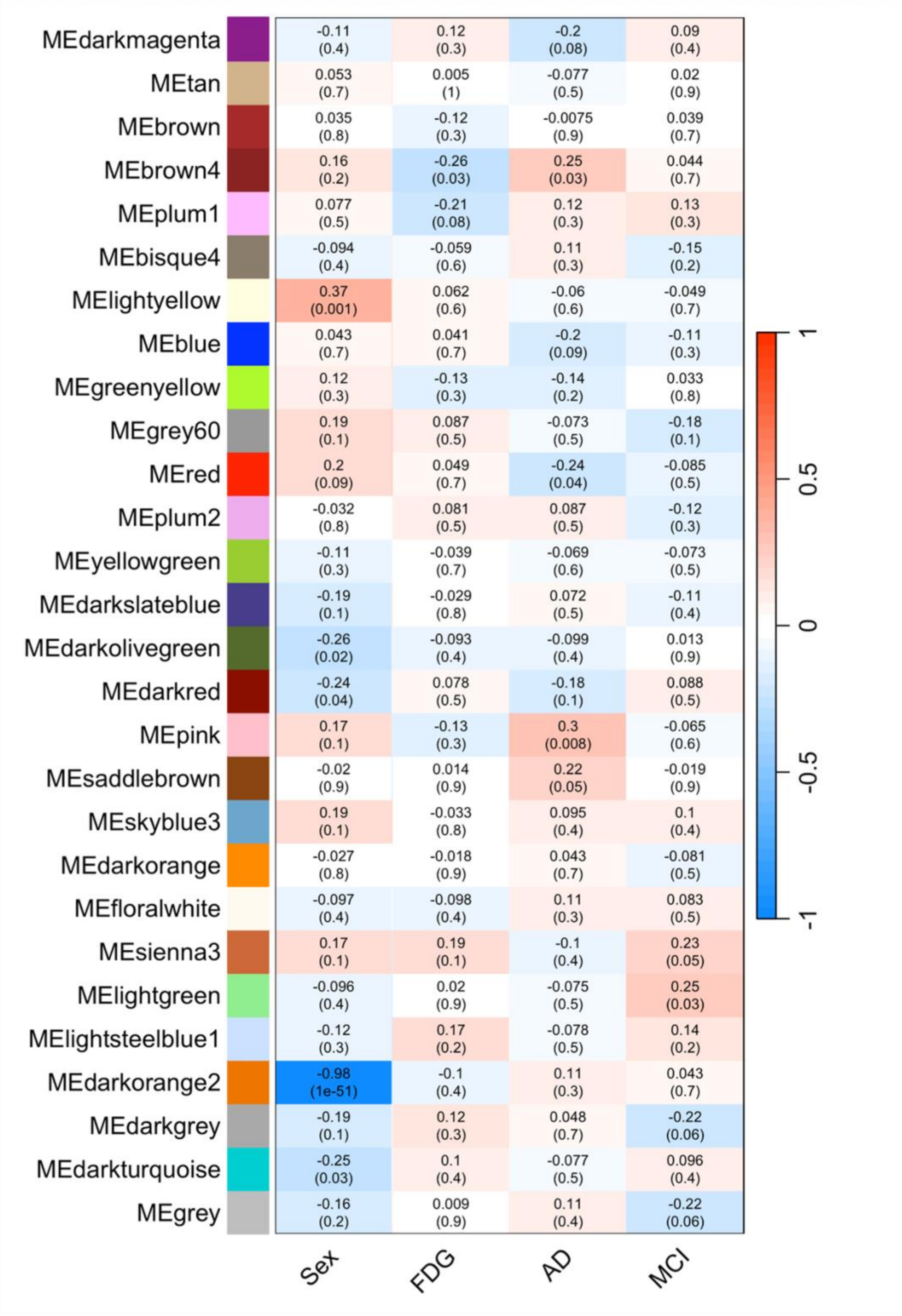


| | Sex | FDG | AD | MCI |
|---|---|---|---|---|
| MEdarkmagenta | -0.11 (0.4) | 0.12 (0.3) | -0.2 (0.08) | 0.09 (0.4) |
| MEtan | 0.053 (0.7) | 0.005 (1) | -0.077 (0.5) | 0.02 (0.9) |
| MEbrown | 0.035 (0.8) | -0.12 (0.3) | -0.0075 (0.9) | 0.039 (0.7) |
| MEbrown4 | 0.16 (0.2) | -0.26 (0.03) | 0.25 (0.03) | 0.044 (0.7) |
| MEplum1 | 0.077 (0.5) | -0.21 (0.08) | 0.12 (0.3) | 0.13 (0.3) |
| MEbisque4 | -0.094 (0.4) | -0.059 (0.6) | 0.11 (0.3) | -0.15 (0.2) |
| MElightyellow | 0.37 (0.001) | 0.062 (0.6) | -0.06 (0.6) | -0.049 (0.7) |
| MEblue | 0.043 (0.7) | 0.041 (0.7) | -0.2 (0.09) | -0.11 (0.3) |
| MEgreenyellow | 0.12 (0.3) | -0.13 (0.3) | -0.14 (0.2) | 0.033 (0.8) |
| MEgrey60 | 0.19 (0.1) | 0.087 (0.5) | -0.073 (0.5) | -0.18 (0.1) |
| MEred | 0.2 (0.09) | 0.049 (0.7) | -0.24 (0.04) | -0.085 (0.5) |
| MEplum2 | -0.032 (0.8) | 0.081 (0.5) | 0.087 (0.5) | -0.12 (0.3) |
| MEyellowgreen | -0.11 (0.3) | -0.039 (0.7) | -0.069 (0.6) | -0.073 (0.5) |
| MEdarkslateblue | -0.19 (0.1) | -0.029 (0.8) | 0.072 (0.5) | -0.11 (0.4) |
| MEdarkolivegreen | -0.26 (0.02) | -0.093 (0.4) | -0.099 (0.4) | 0.013 (0.9) |
| MEdarkred | -0.24 (0.04) | 0.078 (0.5) | -0.18 (0.1) | 0.088 (0.5) |
| MEpink | 0.17 (0.1) | -0.13 (0.3) | 0.3 (0.008) | -0.065 (0.6) |
| MEsaddlebrown | -0.02 (0.9) | 0.014 (0.9) | 0.22 (0.05) | -0.019 (0.9) |
| MEskyblue3 | 0.19 (0.1) | -0.033 (0.8) | 0.095 (0.4) | 0.1 (0.4) |
| MEdarkorange | -0.027 (0.8) | -0.018 (0.9) | 0.043 (0.7) | -0.081 (0.5) |
| MEfloralwhite | -0.097 (0.4) | -0.098 (0.4) | 0.11 (0.3) | 0.083 (0.5) |
| MEsienna3 | 0.17 (0.1) | 0.19 (0.1) | -0.1 (0.4) | 0.23 (0.05) |
| MElightgreen | -0.096 (0.4) | 0.02 (0.9) | -0.075 (0.5) | 0.25 (0.03) |
| MElightsteelblue1 | -0.12 (0.3) | 0.17 (0.2) | -0.078 (0.5) | 0.14 (0.2) |
| MEdarkorange2 | -0.98 (1e-51) | -0.1 (0.4) | 0.11 (0.3) | 0.043 (0.7) |
| MEdarkgrey | -0.19 (0.1) | 0.12 (0.3) | 0.048 (0.7) | -0.22 (0.06) |
| MEdarkturquoise | -0.25 (0.03) | 0.1 (0.4) | -0.077 (0.5) | 0.096 (0.4) |
| MEgrey | -0.16 (0.2) | 0.009 (0.9) | 0.11 (0.4) | -0.22 (0.06) |

**Supplemental Figure S7 – Module-Trait Relationships in Gene Co-expression Network.**
Heatmap of module-trait relationships showing Pearson correlations between WGCNA module eigengenes and clinical traits including sex, cerebral blood flow (CBF), FDG-PET uptake, Alzheimer's disease (AD) status, and mild cognitive impairment (MCI). Each cell represents the strength and direction of the association, quantified by the Pearson correlation coefficient, with statistical significance indicated by the corresponding p-value in parentheses. The color gradient from blue to red represents correlation strength and direction.

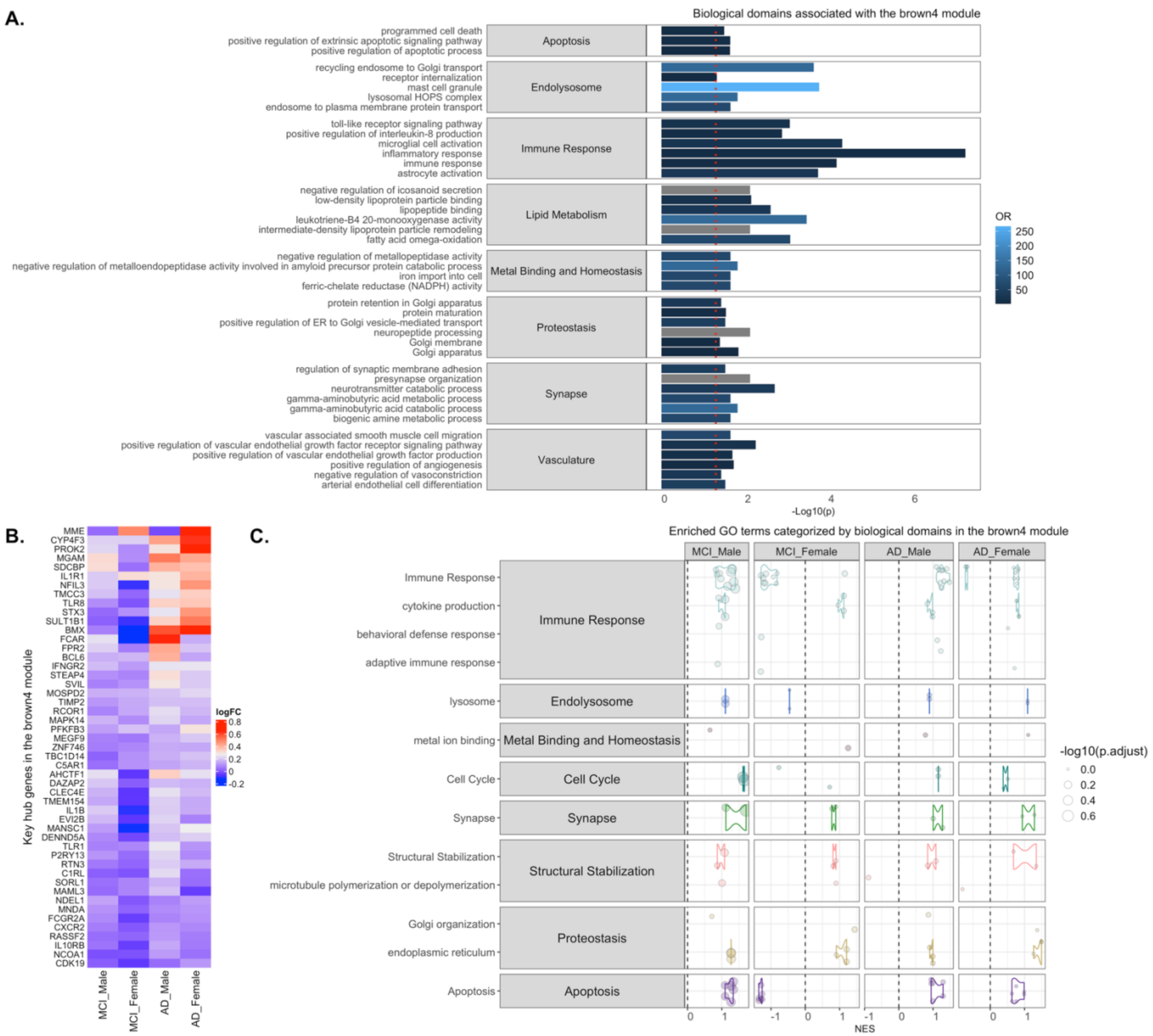


**Supplemental Figure S8 – Functional Characterization of the Brown4 Module.**
Bar plot showing enriched AD-related biological domains and resident GO terms in (**A**) the brown4 module gene set. Each bar represents the strength of enrichment, measured by the odds ratio (OR) and the statistical significance (p-value). Heatmap of log fold change values for (**B**) brown4 module hub genes in MCI and AD cases compared to sex-matched controls. GSEA results for **(C)** brown4 module genes in MCI male, MCI female, AD male, and AD female subgroups. Each point is a GO term within the indicated biological domain, the size of the point is scaled by the GSEA adjusted p-value, and the normalized enrichment scores (NES) indicates if the GO term is up- or downregulated.
