## Supplemental Tables S1-S3 and S5-S7 for "Multiscale Metabolic Covariance Networks Uncover Stage-Specific Biomarker Signatures Across the Alzheimer’s Disease Continuum"

### Supplemental Table S1 – Complete Demographics Stats.

| Stage | Count (n) | | Age (years) | | APOE (n) | | BMI | | | | |
|---|---|---|---|---|---|---|---|---|---|---|---|
| | | | | | (ε2*, ε33, ε4*) | | BMI Category | Male | (n) | Female | (n) |
| CN | M | 54 | M | 76·70 ± 7·26 | M | 7,34,13 | Underweight | NA | 0 | 18·37 ± 0·00 | 1 |
| | | | | | | | Normal | 23·70 ± 1·23 | 18 | 23·01 ± 1·87 | 19 |
| | | | | | | | Overweight | 26·89 ± 1·16 | 31 | 27·96 ± 1·36 | 12 |
| | F | 44 | F | 74·94 ± 6·03 | F | 4,29,11 | Obesity I | 31·51 ± 0·90 | 4 | 32·14 ± 1·61 | 10 |
| | | | | | | | Obesity II | 36·16 ± 0·00 | 1 | NA | 0 |
| | | | | | | | Extreme Obesity | NA | 0 | 49·76 ± 2·02 | 2 |
| EMCI | M | 48 | M | 72·52 ± 6·68 | M | 5,27,16 | Underweight | NA | 0 | 18·39 ± 0·00 | 1 |
| | | | | | | | Normal | 22·92 ± 1·41 | 17 | 21·37 ± 2·02 | 9 |
| | | | | | | | Overweight | 26·92 ± 1·26 | 22 | 27·42 ± 1·51 | 13 |
| | F | 28 | F | 69·72 ± 6·09 | F | 3,15,10 | Obesity I | 32·20 ± 1·62 | 8 | 32·35 ± 0·73 | 5 |
| | | | | | | | Obesity II | 37·34 ± 0·00 | 1 | NA | 0 |
| | | | | | | | Extreme Obesity | NA | 0 | NA | 0 |
| MCI | M | 68 | M | 74·33 ± 6·40 | M | 3,48,17 | Underweight | NA | 0 | 17·83 ± 0·00 | 1 |
| | | | | | | | Normal | 23·38 ± 1·39 | 19 | 22·56 ± 1·76 | 22 |
| | | | | | | | Overweight | 27·42 ± 1·59 | 38 | 27·79 ± 1·55 | 25 |
| | F | 61 | F | 71·56 ± 8·55 | F | 3,36,22 | Obesity I | 31·67 ± 1·46 | 10 | 32·19 ± 1·48 | 10 |
| | | | | | | | Obesity II | NA | 0 | 36·25 ± 0·00 | 1 |
| | | | | | | | Extreme Obesity | 43·20 ± 0·00 | 1 | 40·83 ± 0·97 | 2 |
| LMCI | M | 17 | M | 72·25 ± 7·39 | M | 2,10,5 | Underweight | NA | 0 | NA | 0 |
| | | | | | | | Normal | 24·21 ± 0·47 | 3 | 22·23 ± 1·24 | 10 |
| | | | | | | | Overweight | 27·79 ± 0·98 | 10 | 27·28 ± 1·11 | 10 |
| | F | 25 | F | 72·77 ± 6·14 | F | 1,13,11 | Obesity I | 31·37 ± 1·45 | 4 | 31·73 ± 1·27 | 4 |
| | | | | | | | Obesity II | NA | 0 | NA | 0 |
| | | | | | | | Extreme Obesity | NA | 0 | 45·83 ± 0·00 | 1 |
| AD | M | 49 | M | 74·23 ± 8·53 | M | 1,33,15 | Underweight | 18·10 ± 0·49 | 2 | 16·75 ± 0·00 | 1 |
| | | | | | | | Normal | 22·74 ± 1·91 | 16 | 22·35 ± 1·88 | 21 |
| | | | | | | | Overweight | 27·19 ± 1·66 | 23 | 28·08 ± 1·46 | 9 |
| | F | 37 | F | 74·22 ± 7·64 | F | 0,20,17 | Obesity I | 31·45 ± 1·28 | 5 | 32·29 ± 1·24 | 5 |
| | | | | | | | Obesity II | 37·58 ± 0·00 | 1 | 38·34 ± 0·00 | 1 |
| | | | | | | | Extreme Obesity | 40·50 ± 0·00 | 1 | NA | 0 |

**Supplemental Table S2 – FDG Acquisition and Reconstruction Protocol Used in ADNI 2/3 Cycles.**

| Phase | Manufacturer | Model | Radiotracer (mCi) | Scan Start Time (min) | Acquisition Mode | Duration and Framing | Reconstruction Method | Method Parameters | Grid | FOV | Slice Thickness (mm) | Smoothing |
|---|---|---|---|---|---|---|---|---|---|---|---|---|
| ADNI2 | GE | Discovery STE / VCT | 4·5 – 5·5 | 30 | N/A | 30 min, six x 5min frames | 3D IR | 4 iterations; 20 subsets | 128 x 128 | 256·00 | 3·270 | None |
| | | Discovery ST | | | | | | 4 iterations; 21 subsets | | | | |
| | | Discovery RX | | | | | | | | | | |
| | | Discovery LS | | | | | 3D FORE IR | | | | 4·250 | |
| | | Advance | | | | | | | | | | |
| | Phillips | Gemini TF | | | N/A | 30 min, six x 5min frames | 3D LOR-RAMLA | N/A | 128 x 128 | 256·00 | 2·000 | Sharp |
| | | Gemini / Gemini GXL | | | | | | | | | | |
| | | Allegro | | | | | | | | | | |
| | Siemens | BioGraph mCT | | | LIST-MODE | 30 min, six x 5min frames | OSEM3D | 4 iterations; 12 subsets | 400 x 400 | 407·20 | 2·027 | None |
| | | | | | No LIST-MODE | Two 15min scans | | | | | | |
| | | BioGraph TruePoint (1093/1094) | | | LIST-MODE | 30 min, six x 5min frames | OSEM2D | 4 iterations; 14 subsets | 336 x 336 | 341·04 | 2·027 | None |
| | | | | | No LIST-MODE | Two 15min scans | | | | | | |
| | | | | | LIST-MODE | 30 min, six x 5min frames | OSEM2D | 4 iterations; 16 subsets | 336 x 336 | 341·04 | 2·027 | None |
| | | | | | No LIST-MODE | Two 15min scans | | | | | | |
| | | BioGraph HiRes (1080) | | | LIST-MODE | 30 min, six x 5min frames | OSEM2D | 4 iterations; 14 subsets | 168 x 168 | 341·21 | 2·000 | None |
| | | | | | No LIST-MODE | Two 15min scans | | | | | | |
| ADNI3 | GE | Discovery 600, 610, 690, 710 | 5 ± 10% | 30 | N/A | 30 min, six x 5min frames | 3D IR | 4 iterations; 24 subsets | 192 x 192 | 256·00 | 3·270 | None |
| | | Discovery STE | | | | | | 4 iterations; 20 subsets | 128 x 128 | | | |
| | | | | | | | | 4 iterations; 21 subsets | | | | |
| | | Discovery ST | | | | | 3D FORE IR | | | | 4·250 | |
| | | Discovery LS | | | | | | | | | | |
| | | Advance | | | | | | | | | | |
| | Phillips | Ingenuity TF | | | N/A | 30 min, six x 5min frames | BLOB-OS-TF | N/A | 128 x 128 | 256·00 | 2·000 | Sharp |
| | | Gemini TF | | | | | 3D LOR-RAMLA | | | | | |
| | | Gemini / Gemini GXL | | | | | | | | | | |
| | | Allegro | | | | | | | | | | |
| | Siemens | BioGraph mCT | | | LIST-MODE | 30 min, six x 5min frames | OSEM3D | 4 iterations; 24 subsets | 400 x 400 | 407·20 | 2·027 | None |
| | | | | | No LIST-MODE | Two 15min scans | | | | | | |
| | | BioGraph TruePoint (1093/1094) | | | LIST-MODE | 30 min, six x 5min frames | OSEM3D | 4 iterations; 21 subsets | 336 x 336 | 341·04 | 2·027 | None |
| | | | | | No LIST-MODE | Two 15min scans | | | | | | |
| | | BioGraph HiRes (1080) | | | LIST-MODE | 30 min, six x 5min frames | OSEM2D | 4 iterations; 16 subsets | 168 x 168 | 341·04 | 2·000 | None |
| | | | | | No LIST-MODE | Two 15min scans | | | | | | |

**Supplemental Table S3 – Complete Brain Regions Indices and Names.**

| Region Index – Brain Region Names | | | |
|---|---|---|---|
| 01 - Frontal Pole | 16 - Inferior Temp Gyrus temporooccipit | 31 - Precuneus Cortex | 46 - Planum Temporale |
| 02 - Insular Cortex | 17 - Postcentral Gyrus | 32 - Cuneal Cortex | 47 - Supracalcarine Cortex |
| 03 - Superior Frontal Gyrus | 18 - Sup Parietal Lobule | 33 - Frontal Orbital Cortex | 48 - Occipital Pole |
| 04 - Mid Frontal Gyrus | 19 - Supramarginal Gyrus ant | 34 - Parahippocampal Gyrus ant | 49 - Cerebral White Matter |
| 05 - Inferior Frontal Gyrus pars trian | 20 - Supramarginal Gyrus post | 35 - Parahippocampal Gyrus post | 50 - Lateral Ventricle |
| 06 - Inferior Frontal Gyrus pars operc | 21 - Angular Gyrus | 36 - Lingual Gyrus | 51 - Thalamus |
| 07 - Precentral Gyrus | 22 - Lateral Occipital Cortex Sup | 37 - Temp Fusiform Cortex ant | 52 - Caudate |
| 08 - Temporal Pole | 23 - Lateral Occipital Cortex Inf | 38 - Temp Fusiform Cortex post | 53 - Putamen |
| 09 - Superior Temporal Gyrus ant | 24 - Intracalcarine Cortex | 39 - Temporal Occipital Fusiform Cortex | 54 - Pallidum |
| 10 - Superior Temporal Gyrus post | 25 - Frontal Medial Cortex | 40 - Occipital Fusiform Gyrus | 55 - Brain Stem |
| 11 - Mid Temp Gyrus ant | 26 - Juxtapositional Lobule Cortex | 41 - Frontal Operculum Cortex | 56 - Hippocampus |
| 12 - Mid Temp Gyrus post | 27 - Subcallosal Cortex | 42 - Central Opercular Cortex | 57 - Amygdala |
| 13 - Mid Temp Gyrus temporoocc | 28 - Paracingulate Gyrus | 43 - Parietal Operculum Cortex | 58 - Accumbens |
| 14 - Inferior Temp Gyrus ant | 29 - Cingulate Gyrus ant | 44 - Planum Polare | 59 - Cerebellum |
| 15 - Inferior Temp Gyrus post | 30 - Cingulate Gyrus post | 45 - Heschls Gyrus H1 H2 | |

**Supplemental Table S4 – Literature Review of Brain Regions Functions.**

See attached Excel File called: “S4 – Functional Brain Regions.xlsx”.`

**Supplemental Table S5 – KS Values for the Comparisons for the Network Metrics Relative to the Other Disease Stages.**

| | | Males | | | | Females | | | |
|---|---|---|---|---|---|---|---|---|---|
| Group 1 (Reference) | Group 2 | Cluster Coefficient | Degree Distribution | Positive Node Strength | Negative Node Strength | Cluster Coefficient | Degree Distribution | Positive Node Strength | Negative Node Strength |
| CN | EMCI | 2·08E-02 | 9·82E-02 | 6·06E-02 | 3·34E-01 | 7·75E-01 | 1·60E-04 | 1·53E-01 | 9·05E-01 |
| | MCI | 1·69E-06 | 6·18E-01 | 6·06E-02 | 3·20E-03 | 9·82E-02 | 4·65E-01 | 1·15E-02 | 1·53E-01 |
| | LMCI | 6·12E-07 | 2·88E-05 | 7·34E-09 | 6·12E-07 | 6·18E-03 | 1·60E-04 | 2·08E-02 | 6·06E-02 |
| | AD | 3·34E-01 | 6·18E-01 | 4·65E-01 | 2·31E-01 | 3·20E-03 | 9·79E-01 | 1·53E-01 | 2·08E-02 |
| EMCI | MCI | 9·82E-02 | 6·18E-03 | 7·75E-01 | 3·34E-01 | 6·06E-02 | 1·60E-04 | 1·59E-03 | 7·67E-04 |
| | LMCI | 4·34E-14 | 6·18E-03 | 9·52E-15 | 1·31E-11 | 1·59E-03 | 6·18E-01 | 2·08E-02 | 9·82E-02 |
| | AD | 2·88E-05 | 6·18E-03 | 1·15E-02 | 6·06E-02 | 7·67E-04 | 7·67E-04 | 9·82E-02 | 4·65E-01 |
| MCI | LMCI | 4·12E-16 | 7·20E-08 | 2·02E-15 | 3·31E-12 | 2·88E-05 | 1·69E-06 | 3·56E-04 | 3·56E-04 |
| | AD | 6·12E-07 | 4·65E-01 | 6·18E-03 | 6·18E-03 | 1·69E-06 | 6·18E-01 | 6·91E-05 | 1·60E-04 |
| LMCI | AD | 4·51E-06 | 7·20E-08 | 2·14E-07 | 2·88E-05 | 1·53E-01 | 7·67E-04 | 7·75E-01 | 9·05E-01 |

Yellow boxes represent p<0·05 for KS testing.

**Supplemental Table S6 – Shortest Path Metrics per Disease Stage.**

| Stage | Males | | | | | | Females | | | | | |
|---|---|---|---|---|---|---|---|---|---|---|---|---|
| | Positive Paths Total Length | Negative Paths Total Length | Number of Positive Edges | Number of Negative Edges | Number of Positive Segment Paths | Number of Negative Segment Paths | Positive Paths Total Length | Negative Paths Total Length | Number of Positive Edges | Number of Negative Edges | Number of Positive Segment Paths | Number of Negative Segment Paths |
| CN | 2190·57 | 2000·24 | 361 | 385 | 11216 | 10062 | 2493·89 | 2264·48 | 317 | 351 | 11210 | 10210 |
| EMCI | 2419·82 | 2283·54 | 317 | 351 | 11420 | 10596 | 3268·03 | 3087·02 | 270 | 245 | 11650 | 11172 |
| MCI | 1955·10 | 1768·82 | 362 | 410 | 11096 | 9854 | 2286·68 | 2053·24 | 318 | 340 | 11384 | 10284 |
| LMCI | 4240·36 | 3968·04 | 270 | 255 | 12080 | 11456 | 3460·20 | 3301·62 | 273 | 244 | 11864 | 11340 |
| AD | 2206·13 | 2038·24 | 384 | 414 | 10910 | 9888 | 2640·10 | 2537·43 | 345 | 321 | 11112 | 10512 |

**Supplemental Table S7 – Correlation Between the CCA and the Network and Path Metrics.**

| rho | Density | Degrees | Clustering | Pos Strs | Neg Strs | Pos Paths | Neg Paths | Pos Edges | Neg Edges | Pos Segs | Neg Segs |
|---|---|---|---|---|---|---|---|---|---|---|---|
| Q1SCORE | 0·2810 | 0·2810 | 0·5167 | 0·8360 | 0·6020 | 0·0324 | 0·0582 | 0·4005 | 0·1865 | -0·2239 | -0·0897 |
| Q2SCORE | 0·2770 | 0·2770 | 0·4983 | 0·8779 | 0·5923 | 0·0166 | 0·0510 | 0·4288 | 0·1520 | -0·2959 | -0·0876 |
| Q3SCORE | 0·1841 | 0·1841 | 0·4211 | 0·7017 | 0·4590 | 0·0432 | 0·0635 | 0·2996 | 0·0700 | -0·2629 | -0·0691 |
| Q4SCORE | 0·1809 | 0·1809 | 0·5820 | 0·8199 | 0·5534 | 0·1274 | 0·1519 | 0·3007 | 0·0768 | -0·1354 | 0·0010 |
| Q5SCORE | 0·3573 | 0·3573 | 0·3468 | 0·8010 | 0·5469 | -0·1282 | -0·0965 | 0·4905 | 0·2474 | -0·4215 | -0·2186 |
| Q6SCORE | 0·3436 | 0·3436 | 0·3121 | 0·7816 | 0·5042 | -0·1511 | -0·1133 | 0·4923 | 0·2138 | -0·4385 | -0·2104 |
| Q7SCORE | 0·2375 | 0·2375 | 0·4537 | 0·8293 | 0·5319 | 0·0055 | 0·0399 | 0·3853 | 0·1145 | -0·3255 | -0·0900 |
| Q8SCORE | 0·2462 | 0·2462 | 0·5598 | 0·8612 | 0·6091 | 0·0669 | 0·0934 | 0·3722 | 0·1448 | -0·2096 | -0·0613 |
| Q9SCORE | 0·3787 | 0·3787 | 0·1577 | 0·6959 | 0·3972 | -0·2582 | -0·2206 | 0·5170 | 0·2550 | -0·5433 | -0·3017 |
| Q10SCORE | 0·3754 | 0·3754 | 0·3352 | 0·8151 | 0·6039 | -0·1369 | -0·1041 | 0·5118 | 0·2643 | -0·4662 | -0·2339 |
| Q11SCORE | 0·1494 | 0·1494 | 0·3986 | 0·6834 | 0·3784 | 0·0235 | 0·0514 | 0·2781 | 0·0557 | -0·2455 | -0·0475 |
| Q12SCORE | 0·3913 | 0·3913 | 0·1755 | 0·7227 | 0·4702 | -0·2491 | -0·2114 | 0·5294 | 0·2778 | -0·5650 | -0·3037 |
| Q13SCORE | 0·4696 | 0·4696 | 0·3372 | 0·8143 | 0·6297 | -0·2055 | -0·1868 | 0·5810 | 0·3420 | -0·4766 | -0·3382 |
| CDMEMORY | 0·1281 | 0·1281 | 0·3703 | 0·6443 | 0·3344 | 0·0562 | 0·0895 | 0·2587 | 0·0261 | -0·2001 | 0·0089 |
| CDORIENT | 0·3126 | 0·3126 | 0·4192 | 0·8262 | 0·5787 | -0·0496 | -0·0192 | 0·4445 | 0·1989 | -0·3624 | -0·1585 |
| CDJUDGE | 0·1564 | 0·1564 | 0·3590 | 0·6626 | 0·3453 | 0·0158 | 0·0484 | 0·2905 | 0·0440 | -0·2498 | -0·0352 |
| CDCOMMUN | 0·3132 | 0·3132 | 0·3576 | 0·7982 | 0·5284 | -0·0901 | -0·0539 | 0·4544 | 0·2022 | -0·3980 | -0·1688 |
| CDHOME | 0·2823 | 0·2823 | 0·3946 | 0·7997 | 0·5210 | -0·0475 | -0·0133 | 0·4218 | 0·1713 | -0·3546 | -0·1339 |
| CDCARE | 0·4897 | 0·4897 | 0·2280 | 0·7799 | 0·5769 | -0·2722 | -0·2453 | 0·6076 | 0·3768 | -0·5600 | -0·3739 |
| Visuospatial / Executive | 0·3176 | 0·3176 | 0·2740 | 0·7092 | 0·4802 | -0·1342 | -0·1023 | 0·4417 | 0·1968 | -0·4305 | -0·2080 |
| Naming | 0·3253 | 0·3253 | 0·4015 | 0·7884 | 0·5660 | -0·0636 | -0·0370 | 0·4479 | 0·2005 | -0·3456 | -0·1718 |
| Attention | 0·1764 | 0·1764 | 0·3373 | 0·6614 | 0·3553 | -0·0314 | 0·0029 | 0·3190 | 0·0590 | -0·2878 | -0·0667 |
| Language | 0·2846 | 0·2846 | 0·1889 | 0·5862 | 0·3254 | -0·1644 | -0·1353 | 0·4058 | 0·1900 | -0·3864 | -0·1941 |
| Abstraction | 0·3305 | 0·3305 | 0·1289 | 0·6290 | 0·3124 | -0·2542 | -0·2171 | 0·4740 | 0·1955 | -0·5171 | -0·2706 |
| Delayed Recall | 0·2964 | 0·2964 | 0·4966 | 0·7851 | 0·5790 | 0·0146 | 0·0312 | 0·3960 | 0·1864 | -0·2123 | -0·1239 |
| Orientation | 0·2948 | 0·2948 | 0·4046 | 0·8208 | 0·5424 | -0·0567 | -0·0237 | 0·4363 | 0·1643 | -0·3807 | -0·1546 |

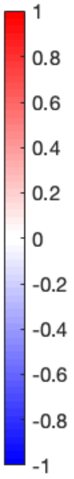


| p-value | Density | Degrees | Clustering | Pos Strs | Neg Strs | Pos Paths | Neg Paths | Pos Edges | Neg Edges | Pos Segs | Neg Segs |
|---|---|---|---|---|---|---|---|---|---|---|---|
| Q1SCORE | 0·4315 | 0·4315 | 0·1262 | 0·0026 | 0·0655 | 0·9292 | 0·8731 | 0·2515 | 0·6059 | 0·5341 | 0·8053 |
| Q2SCORE | 0·4384 | 0·4384 | 0·1427 | 0·0008 | 0·0712 | 0·9637 | 0·8888 | 0·2162 | 0·6751 | 0·4065 | 0·8098 |
| Q3SCORE | 0·6106 | 0·6106 | 0·2255 | 0·0237 | 0·1821 | 0·9058 | 0·8617 | 0·4003 | 0·8476 | 0·4630 | 0·8495 |
| Q4SCORE | 0·6170 | 0·6170 | 0·0775 | 0·0037 | 0·0970 | 0·7257 | 0·6753 | 0·3985 | 0·8329 | 0·7092 | 0·9978 |
| Q5SCORE | 0·3107 | 0·3107 | 0·3262 | 0·0054 | 0·1019 | 0·7241 | 0·7909 | 0·1500 | 0·4907 | 0·2250 | 0·5439 |
| Q6SCORE | 0·3310 | 0·3310 | 0·3799 | 0·0076 | 0·1373 | 0·6769 | 0·7554 | 0·1484 | 0·5530 | 0·2049 | 0·5596 |
| Q7SCORE | 0·5088 | 0·5088 | 0·1878 | 0·0030 | 0·1135 | 0·9881 | 0·9129 | 0·2715 | 0·7528 | 0·3588 | 0·8047 |
| Q8SCORE | 0·4928 | 0·4928 | 0·0924 | 0·0014 | 0·0616 | 0·8542 | 0·7974 | 0·2895 | 0·6899 | 0·5611 | 0·8664 |
| Q9SCORE | 0·2805 | 0·2805 | 0·6635 | 0·0254 | 0·2557 | 0·4713 | 0·5402 | 0·1260 | 0·4771 | 0·1046 | 0·3968 |
| Q10SCORE | 0·2851 | 0·2851 | 0·3438 | 0·0041 | 0·0645 | 0·7060 | 0·7748 | 0·1305 | 0·4606 | 0·1744 | 0·5154 |
| Q11SCORE | 0·6803 | 0·6803 | 0·2538 | 0·0294 | 0·2809 | 0·9486 | 0·8879 | 0·4366 | 0·8786 | 0·4943 | 0·8964 |
| Q12SCORE | 0·2635 | 0·2635 | 0·6278 | 0·0182 | 0·1703 | 0·4877 | 0·5577 | 0·1156 | 0·4370 | 0·0888 | 0·3936 |
| Q13SCORE | 0·1708 | 0·1708 | 0·3407 | 0·0041 | 0·0511 | 0·5689 | 0·6053 | 0·0782 | 0·3333 | 0·1637 | 0·3392 |
| CDMEMORY | 0·7244 | 0·7244 | 0·2922 | 0·0444 | 0·3450 | 0·8774 | 0·8057 | 0·4704 | 0·9430 | 0·5794 | 0·9805 |
| CDORIENT | 0·3792 | 0·3792 | 0·2278 | 0·0032 | 0·0796 | 0·8917 | 0·9580 | 0·1981 | 0·5818 | 0·3034 | 0·6619 |
| CDJUDGE | 0·6662 | 0·6662 | 0·3083 | 0·0368 | 0·3284 | 0·9655 | 0·8943 | 0·4155 | 0·9040 | 0·4863 | 0·9231 |
| CDCOMMUN | 0·3782 | 0·3782 | 0·3103 | 0·0056 | 0·1164 | 0·8044 | 0·8825 | 0·1871 | 0·5754 | 0·2547 | 0·6411 |
| CDHOME | 0·4294 | 0·4294 | 0·2591 | 0·0055 | 0·1225 | 0·8964 | 0·9710 | 0·2247 | 0·6362 | 0·3147 | 0·7123 |
| CDCARE | 0·1508 | 0·1508 | 0·5265 | 0·0078 | 0·0808 | 0·4468 | 0·4946 | 0·0624 | 0·2832 | 0·0923 | 0·2872 |
| Visuospatial / Executive | 0·3713 | 0·3713 | 0·4435 | 0·0216 | 0·1601 | 0·7117 | 0·7785 | 0·2012 | 0·5858 | 0·2142 | 0·5642 |
| Naming | 0·3591 | 0·3591 | 0·2501 | 0·0067 | 0·0881 | 0·8615 | 0·9191 | 0·1942 | 0·5786 | 0·3281 | 0·6350 |
| Attention | 0·6258 | 0·6258 | 0·3406 | 0·0373 | 0·3137 | 0·9314 | 0·9936 | 0·3690 | 0·8715 | 0·4200 | 0·8548 |
| Language | 0·4255 | 0·4255 | 0·6011 | 0·0749 | 0·3589 | 0·6499 | 0·7093 | 0·2447 | 0·5990 | 0·2700 | 0·5910 |
| Abstraction | 0·3510 | 0·3510 | 0·7226 | 0·0514 | 0·3795 | 0·4784 | 0·5469 | 0·1663 | 0·5882 | 0·1259 | 0·4495 |
| Delayed Recall | 0·4057 | 0·4057 | 0·1443 | 0·0071 | 0·0795 | 0·9681 | 0·9319 | 0·2573 | 0·6062 | 0·5559 | 0·7332 |
| Orientation | 0·4083 | 0·4083 | 0·2461 | 0·0036 | 0·1053 | 0·8765 | 0·9482 | 0·2075 | 0·6501 | 0·2778 | 0·6698 |

**Note**: The first table presents the correlation coefficient between each test and each metric. The subsequent table provides the p-value for each comparison, indicating that only the positive path lengths exhibit significant variations (The green box represents p<0·05). This suggests that the CCA scores alone are insufficient to capture the alterations in the metabolic covariance matrices or the changes in the path lengths.
